# A hierarchy of functional states in working memory

**DOI:** 10.1101/2020.04.16.044511

**Authors:** Paul S. Muhle-Karbe, Nicholas E. Myers, Mark G. Stokes

## Abstract

Extensive research has examined how information is maintained in working memory (WM), but it remains unknown how WM is used to guide behaviour. We addressed this question by combining human electrophysiology with pattern analyses, cognitive modelling, and a task requiring maintenance of two WM items and priority shifts between them. This enabled us to discern neural states coding for immediately and prospectively task-relevant items, and to examine their contribution to WM-based decisions. We identified two qualitatively different states: a functionally active state encoded only immediately task-relevant items and closely tracked the quality of evidence integration on the current trial. In contrast, prospectively relevant items were encoded in a functionally latent state that did not engage with ongoing processing but tracked memory precision at longer time scales. These results delineate a hierarchy of functional states, whereby latent memories supporting general maintenance are transformed into active decision-circuits to guide flexible behaviour.

## Introduction

Working memory (WM) refers to the ability to maintain and manipulate information that is no longer available in the environment. It provides a flexible mental workspace that scaffolds many higher cognitive functions such as planning, reasoning, or cognitive control.^1,2^ As a consequence, a longstanding and central theme in cognitive neuroscience has been to delineate neural mechanisms that underpin WM.^3–6^ This research has greatly advanced our understanding of mechanisms that enable the brain to maintain WM contents over delay periods, e.g., via persistent activity,^7–9^ neuronal oscillations,^10,11^ or activity-silent brain sates.^12–14^ At the same time, however, this work has so far neglected a key aspect of WM, namely how its contents are used to guide flexible behaviour.

Existing evidence suggests that the use of memories is distinct from the mere maintenance of information.^15^ For instance, patients with frontal lobe damage sometimes exhibit a phenomenon termed goal neglect, wherein instructed task components are disregarded during performance, even though they have been understood and remembered.^16,17^ Studies in healthy humans furthermore suggest that WM items that will guide the next decision can bias perception toward memory-matching stimuli, while other concurrently held items do not affect ongoing cognition.^18^ Likewise, in neuroimaging studies, immediately task-relevant WM items can be robustly decoded from brain activity patterns, while other concurrently held items typically exhibit little or no decodability.^19–23^ Such findings are commonly thought to reflect the effects of attention, which may prioritise a single item in WM by amplifying the corresponding neural patterns.^24–27^ This view echoes the tenet of models conceiving WM storage as the distribution of limited cognitive resources. From this perspective, selecting an item for guiding behaviour focusses resources, boosting the strength of the selected item, while impeding the strength of other concurrently held items.

An alternative proposal states that WM-guided behaviour that does not rely on attentional selection alone, but also entails a reconfiguration of representational formats from a purely mnemonic state into an action-ready state that is tuned for efficient task-dependent decision making.^15,28^ From this perspective, WM representations should exhibit qualitatively different functional attributes depending on their momentary task relevance. Items that are immediately relevant for guiding behaviour should be encoded in a *functionally active* state that is optimised for efficient task-dependent readout. By contrast, items that are not immediately task-relevant, but will become relevant later on, should be held in a *functionally latent* state that enables precise maintenance, but does not modulate cognition until it becomes relevant for guiding behaviour.

In the current study, we tested this view using electroencephalography (EEG) and a novel task that allowed us to directly compare these two putative functional states. Participants were asked to hold two items in WM for an extended period of time, though at any given moment only one item was immediately task-relevant (cued item), while the other item was maintained for later use (uncued item). We leveraged a combination of multivariate decoding analyses and cognitive modelling to recover representations of cued and uncued WM items, and to link variance in their decoding strength with variance in performance. Consistent with our proposal, we find that cued and uncued WM items are encoded in largely dissociable neural patterns. Neural patterns coding for uncued items reflect a functionally latent mnemonic space that tracks the precision of maintenance at longer time scales but is unrelated to trial-wise variance in performance. By contrast, the unique pattern component coding for cued items reflects a functionally active state that closely tracks trial-wise variance in the quality of evidence integration for WM-based decisions. Functional states are moreover highly flexible and can be dynamically shifted between items without degrading their precision. These findings suggest a hierarchy of functional states in WM, wherein functionally latent memories that support general maintenance can be transformed into functionally active decision circuits to guide flexible behaviour.

## Results

### Behaviour

50 healthy adults participated in two experiments (20 participants in experiment 1 and 30 participants in experiment 2) while undergoing EEG recordings. Experiment 2 served as a conceptual replication of experiment 1. Data were also pooled across experiments for several analyses to maximise statistical power (Materials and Methods). Participants performed a WM-based decision-making task that required maintenance of two items for a block of 16 trials, and flexible priority shifts between those items on a trial-wise basis. At the beginning of each block, participants were shown two orientated bars in blue and yellow colour, which served as WM items. During the block, each trial started with the presentation of an auditory cue that signalled which of the two WM items should be used as a reference for a forthcoming perceptual decision (Figure 1). Accordingly, on any given trial, only one of the two WM items was task-relevant (cued item), while the other item had to be maintained for use later in the block (uncued item). The cue was followed by a 700ms delay period until a randomly oriented Gabor patch was presented as target. In experiment 1, the target was presented centrally on the screen, while it was presented peripherally in experiment 2 (Materials and Methods). Participants were given a maximum time window of 4s to classify the target via right-hand or left-hand button press as being tilted clockwise or counter-clockwise relative to the cued memory item. Importantly, participants saw the memory items only during the encoding phase at the block start and did not receive trial-wise response feedback. Accordingly, to solve the task, they had to maintain precise representations of both WM items throughout the block.

**Figure1:**
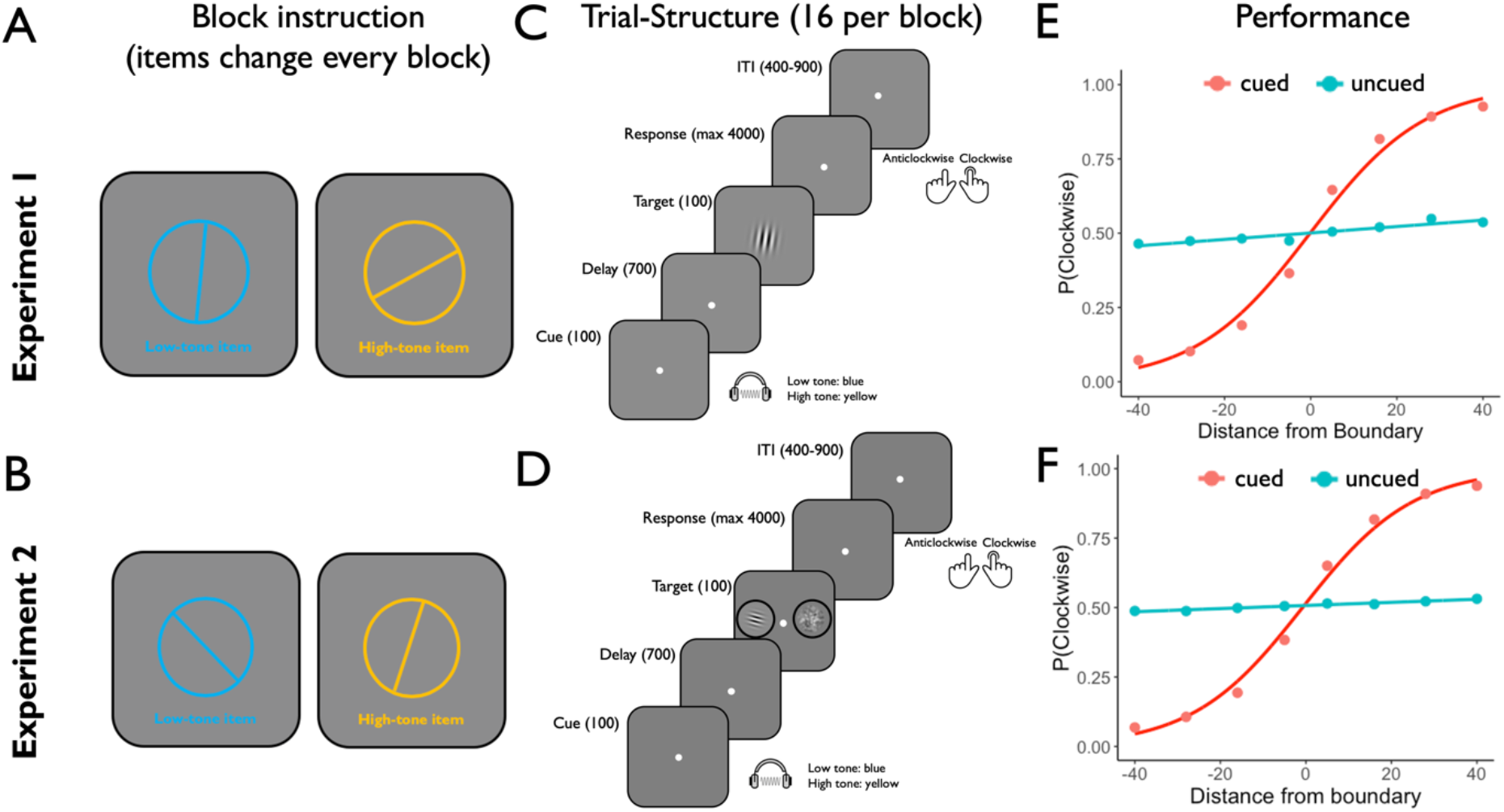
Illustration of the task design and behavioural results of experiment 1 (upper panel) and experiment 2 (bottom panel). **A,B** At the beginning of each block, two randomly oriented bars were shown successively in blue and yellow colour. These two items served as WM items for the remainder of the block and were associated with a high-pitch tone and a low-pitch tone respectively (mapping counterbalanced across subjects). **C,D** During the block, each trial started with the presentation of an auditory cue that signals which WM item should be used as decision criterion on the current trial (cued item), while the other item had to be maintained for later use in the block (uncued item). After a brief delay, a randomly oriented gabor patch was presented as target, and participants were required to indicate if the target was oriented clockwise or anticlockwise relative to the active WM item. In experiment 1, the target was shown centrally on the screen (C). In experiment 2, the target was shown peripherally on the left or right side of the fixation dot, while a noise patch was presented on the other side (D). The noise patch matched the target in luminance, size, contrast, and eccentricity. Note that the location of the target was indicated to participants at the beginning of each block. **E,F** Probability of clockwise responses as a function of the angular difference between target stimulus and the currently cued WM item (shown in red) and from the currently uncued WM item (shown in blue). Data are shown in dots and the lines represent a fitted binomial cumulative distribution function. Responses were strongly modulated by the angular distance between the orientation of the target and the orientation of the cued item, whereas the distance between the target and uncued item had only minimal impact on performance (see Material and Methods and Results for details).

Participants were able to perform the task well above chance (experiment 1: accuracy = 79.5%, reaction time = 573ms; experiment 2: accuracy = 80.1% reaction time = 679ms). As shown in Figure 1, performance was strongly modulated by the angular distance between the target orientation and the orientation of the cued WM item. Reaction time (RT) decreased with greater distances (experiment 1: t_19_ = 10.078, p < 0.001; experiment 2: t_29_ = 10.462, p < 0.001), while accuracy increased (experiment 1: t_19_ = 12.677, p < 0.001; experiment 2: t_29_ = 18.426, p < 0.001). By contrast, the angular distance between the target and the uncued WM item had no reliable impact on RT (experiment 1: t_19_ = 2.49, p = 0.069; experiment 2: t_29_ =2.125, p = 0.103), and only a modest effect on accuracy (experiment 1: t_19_ = 3.12, p = 0.033; experiment 2: t_29_ = 4.18, p = 0.050). In addition, we observed a reliable effect of priority shifts, whereby performance was slower (experiment 1: t_19_ = 7.097, p < 0.001; experiment 2: t_29_ = 12.864, p < 0.001) and more error prone (experiment 1: t_19_ = −2.704, p = 0.014; experiment 2: t_29_ = −2.711, p = 0.011) on trials that incurred a shift in priority between the two WM items, relative to the previous trial, than on trials in which the item priority repeated (see supplementary Figure 1). We also found that accuracy was stable across different experimental blocks (experiment 1: t = 1.214, p = 0.240; experiment 2: t_29_ = 0.080, p = 0.937), but declined continuously across trials within blocks (experiment 1: t_19_ = −13.841, p < 0.001; experiment 2: t_29_ = −6.242, p < 0.001). Conversely, RT decreased with increasing block numbers (experiment 1: t = −7.227, p < 0.001; experiment 2: t = −7.129, p < 0.001), but was stable within blocks except for the first block trial (experiment 1: t_19_ = −0.432, p = 0.671, experiment 2: t_29_ = −6.242, p < 0.001; supplementary Figure 2).

### Time-resolved decoding of task-variables

We conducted a series of multivariate pattern analyses to characterise the neural representations that underpin performance in our task. In a first step, we conducted a time-resolved decoding analysis to reveal the time courses within which three task variables were explicitly encoded in EEG sensor activity: 1) the orientation of the target stimulus, 2) the orientation of the WM item, separately for cued and uncued conditions, and 3) a decision variable that was calculated as the absolute distance between the orientation of the target and the orientation of the WM item, again separately for cued and uncued conditions. To recover information about these variables from EEG sensor activity with high temporal resolution, we computed Mahalanobis distances between the patterns of sensor activity that were evoked by different stimulus orientations, and measured the extent to which these distances reflected the underlying circular orientation space or linear decision variable space (see Figure 2 for illustration and Materials and Methods for details).

**Figure 2:**
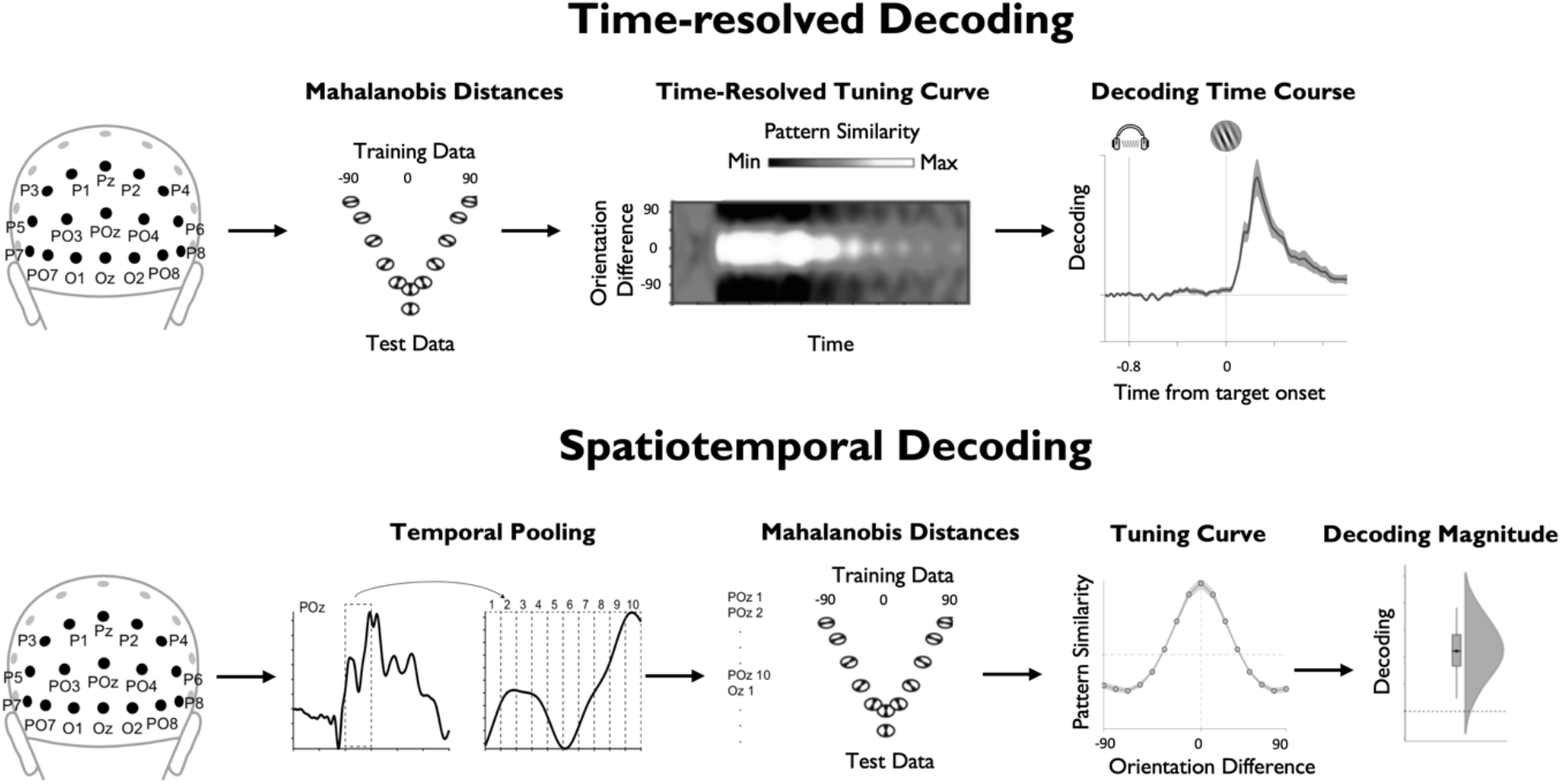
Illustration of time-resolved and spatiotemporal approaches that were used for the decoding of task variables. The upper panel illustrates the time-resolved decoding approach. Here, for each training set and time point, Mahalanobis distances are computed between activation patterns of posterior sensors of the test data and the training data (binned based on their orientation relative to the test data). After repeating this procedure for every training set and time point, the resulting distances were sign-reversed, so that positive scores reflect pattern similarity, and centred around the mean voltage across sensors for each time point. This was used to reconstruct a time-resolved population tuning curve that expresses the extent to which pattern similarity reflects the similarity of the underlying angular orientation space across time. We estimated the height of this tuning curve for each time point by convolving it with a cosine function (see Materials and Methods for details). The resulting vector served as index of time-resolved decoding accuracy. The bottom panel illustrates the spatiotemporal decoding approach. Here, the data entering the decoders were pooled over multiple time points, so as to exploit not only information that is encoded in spatial activation patterns, but also information that is encoded in their temporal structure. We focused on the time window from 100-400ms after target onset and treated individual trials as discrete events. Tuning curves were estimated based on mean-centred and sign-reversed Mahalanobis distances, as in the time-resolved approach, yielding a single index of decoding strength for each trial as final output.

Information about the orientation of the target stimulus started to be encoded in EEG sensor patterns briefly after target onset and decoding remained significant for the remainder of the selected time window (Figure 3; experiment 1: from 72ms after the target, cluster-corrected p < 0.0001; experiment 2: from 68ms after the target, cluster corrected p < 0.0001). Cued and uncued WM items were both decodable, but with clear differences in their respective time courses. Decoding of the cued item started to rise during delay period and was further magnified after target onset (experiment 1: first cluster: 208-376ms after the cue; cluster corrected p = 0.011; second cluster = 400-2000ms after the cue, cluster corrected p < .00001; experiment 2: 420-2000ms after the cue, cluster-corrected p < 0.0001). In contrast, the uncued item was not decodable during the delay period, but there was a time window of significant decoding after the onset of the target stimulus (experiment 1: 192-528ms after the target, cluster corrected p = 0.0005, experiment 2: 272-316ms after the target, cluster corrected p = 0.048). Slightly later during the target period, we could decode a decision variable for the cued item (experiment 1: 244-1000ms after the target, cluster corrected p = 0.0002; experiment 2: 388-1000ms after the target, cluster corrected p = 0.001), whereas no decision variable was decodable for the uncued item. Finally, for all three task variables, we also validated that eye-movements could not explain the pattern of EEG decoding results (supplementary Figures 4 and 5).

**Figure 3:**
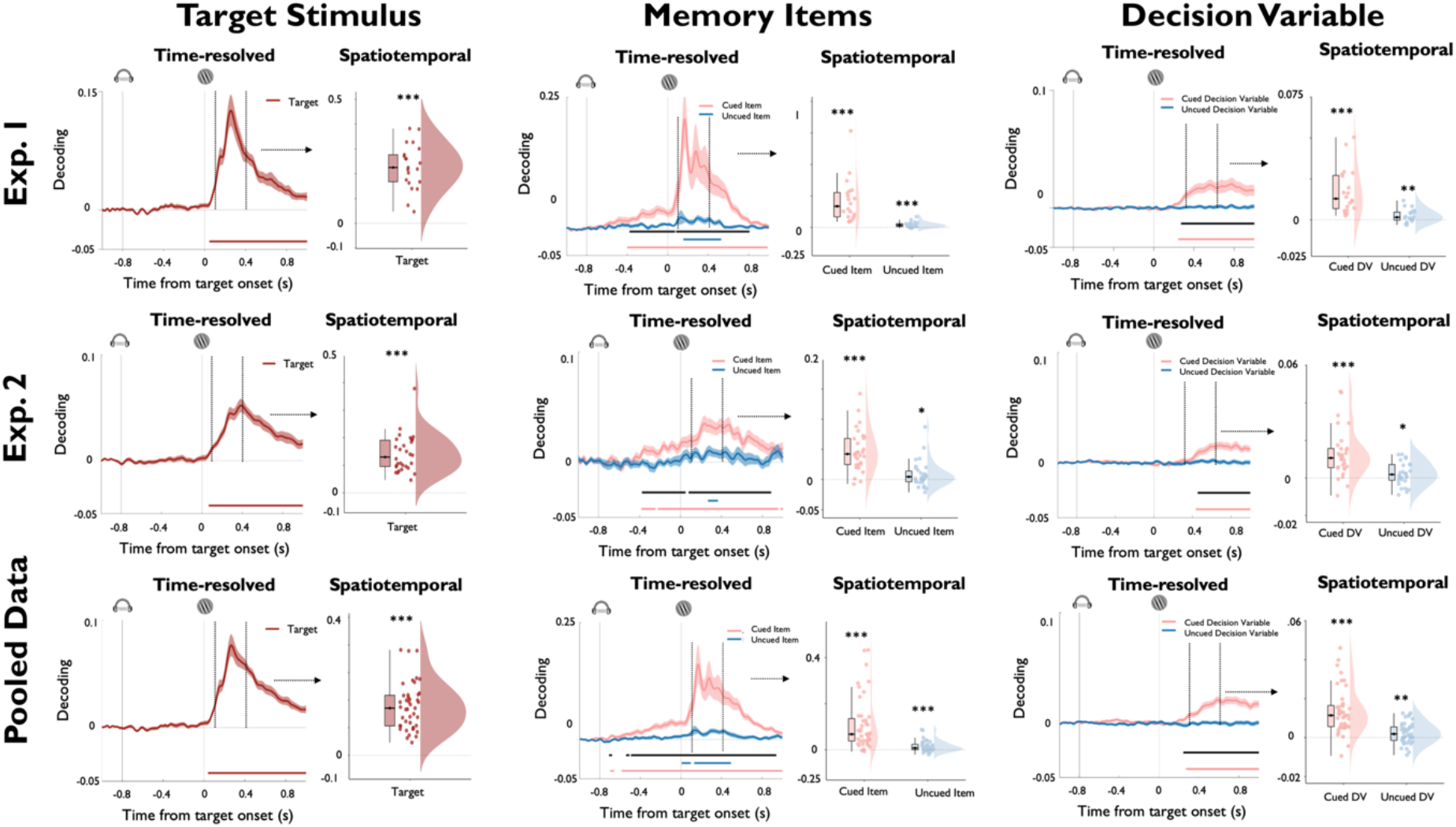
Time-resolved and spatiotemporal decoding of task variables for experiment 1 (first row), experiment 2 (second row) and the pooled data across experiments (third row). Panels on the left display the decoding results for the orientation of the target stimulus, middle panels display the decoding results for the orientation for the cued and uncued WM items, and panels on the right display the decoding results for the absolute value of the decision variable, calculated separately with respect to the cued and uncued WM item. For each experiment and task variable, plots on the left display time-resolved decoding results from 200ms before cue onset until 800ms after target onset. Coloured lines represent cluster-corrected time-periods within which decoding was significantly greater than chance. Black lines in the middle and bottom rows indicate cluster-corrected time periods within which decoding strength of the cued and uncued WM items differed significantly. Shading indicates standard error of the mean. Plots on the right display results of the spatiotemporal decoding approach where data entered into the decoder were pooled within a time window from 100-400ms following target onset (see text for details). Boxplots: centre lines indicate the median, the box outlines indicate the 25^th^ and 75^th^ percentiles, and the whiskers indicate 1.5 times the interquartile range. Significance values: *** p < 0.005; ** p< 0.01; * p < 0.05.

### Spatiotemporal Decoding

In a second step, we employed a complementary analysis approach to decode the same task variables from longer time-windows of stimulus-evoked brain activity. This approach exploits the dynamic temporal structure of event-related potentials by pooling multivariate information in time, thus capturing not only information encoded in spatial activation patterns, but also information encoded in the temporal unfolding of these patterns (Figure 2 and Materials and Methods). Previous work has shown that such pooling over adjacent time points within trials can increase decoding sensitivity at the expense of temporal precision.^29,30^ In keeping with these studies, we pooled data points over a time period from100-400ms after target onset and treated individual trials as discrete events. As shown in Figure 3, this approach clearly enhanced decoding sensitivity. Specifically, we could reliably decode the orientation of the target (experiment 1: t_19_ = 11.585, p < 0.001; experiment 2: t_29_ = 11.301, p < 0.001), the orientation of the cued item (experiment 1: t_19_ = 5.599, p < 0.001; experiment 2: t_29_ = 8.400, p < 0.001) and the orientation of the uncued item (experiment 1: t_19_ = 4.232, p < 0.001; experiment 2: t_29_ = 2.253, p = 0.016). Interestingly, using the spatiotemporal approach, we could not only decode a decision variable for the cued item (experiment 1: t_19_ = 5.922, p < 0.001; experiment 2: t_29_ = 5.233, p < 0.001), but also for the uncued item (experiment 1: t_19_ = 2.627, p = 0.008; experiment 2: t_29_ = 1.889, p = 0.034).

### Regression analyses

#### (1) Active, but not latent, WM states predict trial-wise variance in task performance

We next conducted a series of regression analyses to establish the relevance of the decoded memory signals for task performance and test our hypothesis that cued and uncued items are encoded in qualitatively different functional states. To reiterate, we expected that the cued item would be encoded in a decision-oriented state that directly predicts variance in performance, whereas the uncued item should be held in a functionally latent state that has minimal impact on current performance. To test this prediction, we regressed log-transformed reaction time (log-RT) and accuracy against the trial-wise decoding strength of cued and uncued WM items using the spatiotemporal classifier (see Materials and Methods). As shown in Figure 4, we observed that higher trial-wise decoding of the cued item reliably predicted faster log-RT (experiment 1: t_19_ = −4.362, p < 0.001, Bayes Factor in favour of the alternative hypothesis (BF_H1_) = 190.357; experiment 2: t_29_ = −2.895, p = 0.004, BF_H1_ = 11.950; pooled data: t_42_ = −4.637, p < 0.001, BF_H1_ = 1271.980), but had no reliable impact on accuracy (experiment 1: t_19_ = 2.207, p = 0.020, BF_H1_ = 3.239; experiment 2: t_29_ = 1.504, p = 0.072, BF_H1_ = 0.985; pooled data: t_42_ = 1.596, p = 0.059, BF_H1_ = 0.994). By contrast, trial-wise variance in decoding of the uncued item neither predicted log-RT (experiment 1: t_19_ = 0.622, p = 0.271, Bayes Factor in favour of the null hypothesis (BF_H0_) = 2.515; experiment 2: t_29_ = −0.713, p = 0.759, BF_H0_ = 8.172; pooled data: t_42_ = 0.391, p = 0.349, BF_H0_ = 4.356) nor accuracy (experiment 1: t_19_ = −0.410, p = 0.343, BF_H0_ = 3.072; experiment 2: t_29_ = 1.186, p = 0.877, BF_H0_ = 10.324; pooled data: t_42_ = 0.986, p = 0.835, BF_H0_ = 11.199) with Bayes Factors indicating substantial evidence in favour of the respective null hypotheses. Regression weights of the cued and the uncued items differed significantly in the prediction of log-RT (experiment 1: t_19_ = −2.452, p = 0.012, BF_H1_ = 4.903; experiment 2: t_29_ = −1.360, p = 0.092, BF_H1_ = 0.805; pooled data: t_42_ = −2.669, p = 0.005, BF_H1_ = 7.413). Together, these results support our prediction that cued items are encoded in a functionally active state that guides WM-based decisions, whereas uncued items are held in a functionally latent state that has minimal impact on ongoing cognition.

**Figure 4:**
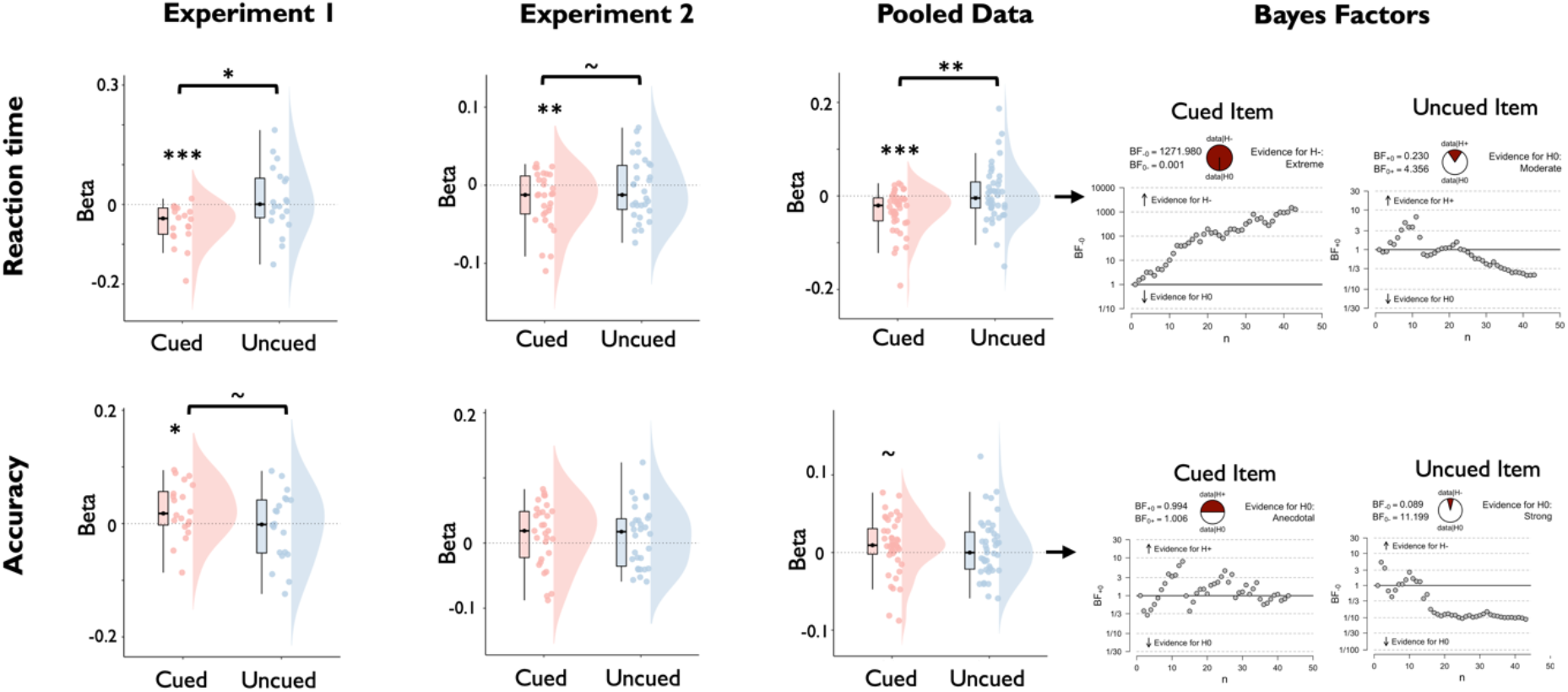
Regression of task performance based on trial-wise variance in decoding strength of the cued and uncued WM items, separately for experiment 1 and experiment 2, and the pooled data across experiments. The top row displays the distribution of regression weights across participants for the prediction of reaction time and the bottom row displays the for the prediction of accuracy. Boxplots: centre lines indicate the median, the box outlines indicate the 25^th^ and 75^th^ percentiles, and the whiskers indicate 1.5 times the interquartile range. Significance values: *** p < 0.005; ** p< 0.01; * p < 0.05. Plots on the right display sequential Bayes Factors for the respective regression weights (based on the pooled data set).

#### (2) Functionally active and latent WM states both track memory precision over longer time scales

After establishing that the decoding strength of uncued items is unrelated to performance on the current trial, we next tested the extent to which these signals nonetheless capture behaviourally relevant variance by examining their contribution to general maintenance. Although a strong representation of the uncued item is not advantageous on the current trial, it is nonetheless crucial for performance on other trials in the same block when priorities differ, and the same item becomes task-relevant. Therefore, maintaining an accurate representation of the uncued WM item reflects a minimal requirement for solving the task as a whole. Accordingly, we reasoned that the overall quality with which a WM item is encoded throughout the block when it is uncued may track performance in the same block on those trials when the same item is cued. We tested this assumption by regressing log-RT and accuracy against the decoding strength of the two memory items averaged over the remaining trials of the block (see Figure 10 and Materials and Methods). Assuming that block-wise decoding tracks differences in the general quality with which items are encoded throughout the block, we reasoned that the strength of both cued and uncued items should scale positively with performance.

As shown in Figure 6, accuracy was indeed predicted by the block-wise decoding strength of the cued item (experiment 1: t_19_ = 3.034, p < 0.003, BF_H1_ = 14.020; experiment 2: t_29_ = 2.526, p = 0.009, BF_H1_ = 5.633; pooled data: t_42_ = 3.797, p < 0.001, BF_H1_ = 177.778) and the uncued item (experiment 1: t_19_ = 2.005, p < 0.030, BF_H1_ = 2.341; experiment 2: t_29_ = 1.656, p = 0.054, BF_H1_ = 1.233; pooled data: t_42_ = −2.318, p < 0.013, BF_H1_ = 5.725). Differences between the regression weights for the cued and the uncued items were nonsignificant, with Bayes Factors providing evidence in favour of the null hypothesis (experiment 1: t_19_ = 0.045, p = 0.964, BF_H0_ = 4.300; experiment 2: t_29_ = 0.826, p = 0.415, BF_H0_ = 3.760; pooled data: t_42_ = 0.514, p = 695, BF_H0_ = 5.796). In addition, block-wise decoding of the cued item also predicted log-RT (experiment 1: t_19_ = −4.093, p < 0.001, BF_H1_ = 111.097; experiment 2: t_29_ = −1.812, p = 0.040, BF_H1_ = 1.575; pooled data: t_42_ = −4.253, p < 0.001, BF_H1_ = 155.046), which was not the case for the uncued item (all p > 0.299; all BF_H0_ > 3.253). Together, these results show that even though functionally latent WM states do not engage with ongoing processing, they nonetheless support performance at longer timescales.

**Figure 6:**
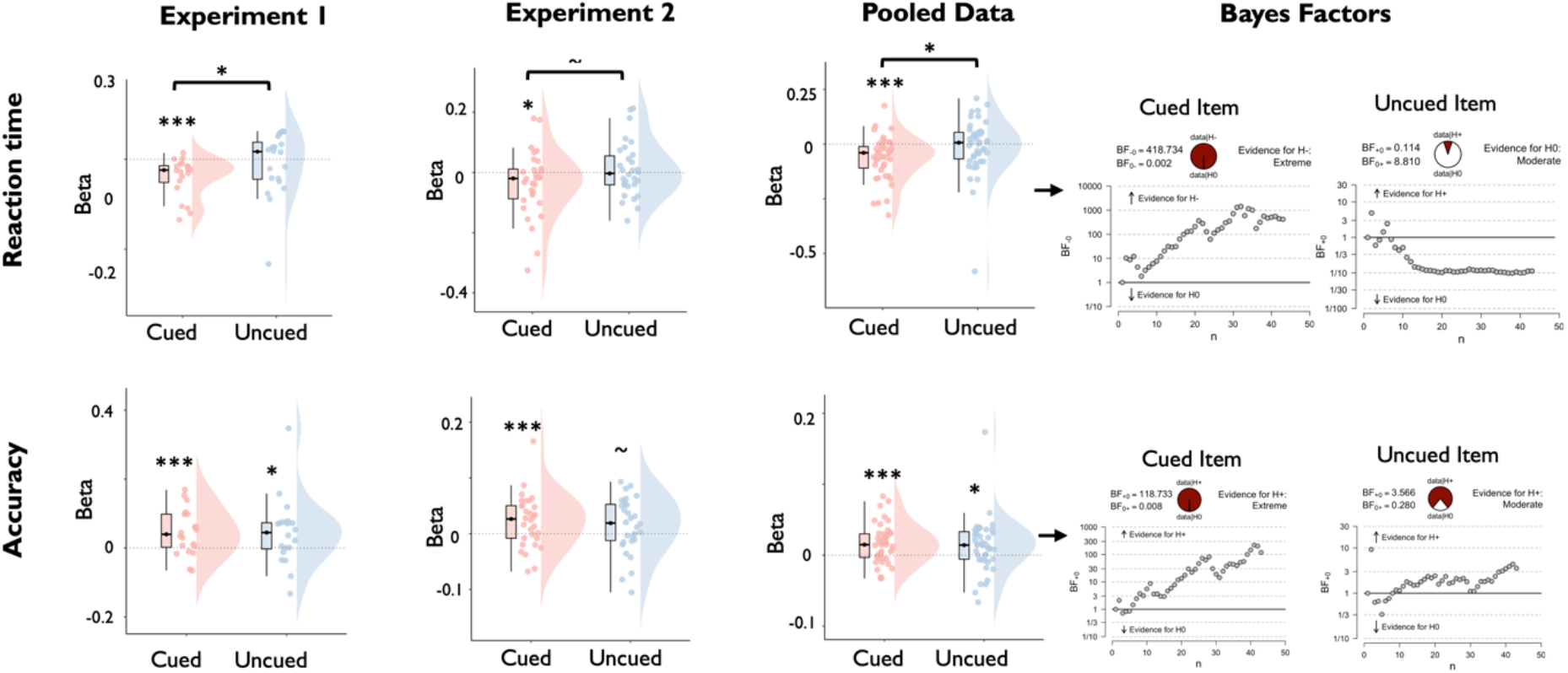
Regression of task performance based on block-wise decoding accuracy of the cued and uncued WM items, separately for experiment 1, experiment 2, and the pooled data across experiments. The top row displays the distribution of regression weights across participants for the prediction of reaction time and the bottom row displays the same for the prediction of accuracy. Boxplots: centre lines indicate the median, the box outlines indicate the 25^th^ and 75^th^ percentiles, and the whiskers indicate 1.5 times the interquartile range. Significance values: *** p < 0.005; ** p< 0.01; * p < 0.05. Plots on the right display sequential Bayes Factors for the respective regression weights (based on the pooled data set).

#### (3) Similarity to the functionally active state predicts interference from uncued items

The previous sections established that uncued WM items are encoded in a format that supports maintenance but has minimal impact on ongoing processing. Nonetheless, we had observed two behavioural signatures of interference between WM items: a subtle but reliable effect of the angular distance between the uncued item and the target stimulus on decision accuracy, and a performance cost on trials demanding a priority shift between WM items. Given our previous finding that the strength of functionally active, but not functionally latent, WM states tracks fluctuations in performance, we reasoned that interference may reflect the extent to which uncued WM items are encoded in patterns resembling their functionally active state. To test this idea, we repeated the trial-wise regression analyses used above with a cross-item decoder that was trained on data sorted by the cued item and tested on data sorted by the uncued item.

The cross-item decoder successfully classified uncued WM items with above-chance accuracy (supplementary Figure 6; experiment 1: t_19_ = 4.719, p < 0.001; experiment 2: t_29_ = 2.751, p = .010). Decoding magnitude was enhanced in comparison to the regular decoder of the uncued item in experiment 1 (t_19_ = 2.74, p = 0.013), but not in experiment 2 (t_29_ = 0.423, p = 0.675). Critically, higher trial-wise cross-item decoding indeed predicted slower log-RT (Figure 7; experiment 1: t_19_ = 3.497, p = 0.001, BF_H1_ = 34.074; experiment 2: t_29_ = 1.996, p = 0.028, BF_H1_ = 2.135; pooled data: t42.= 3.685, p < 0.001, BF_H1_ = 87.960), whereas accuracy was not reliably affected (experiment 1: t_19_ = −1.980, p = 0.031, BF_H1_ = 2.250; experiment 2: t_29_ = −0.772, p = 0.223, BF_H0_ = 2.545; pooled data: t42. = 1.619, p = 0.056, BF_H1_ = 1.032). Notably, the regression weights for log-RT were significantly larger than those of the regular decoder of the uncued item (experiment 1: t_19_ = 3.037, p = 0.003, BF_H1_ = 14.102, experiment 2: t_29_ = 1.907, p = 0.033, BF_H1_ = 1.837; pooled data: t_42_ = 3.201, p = 0.001, BF = 25.515). We tested if the difference between the two decoders could be attributed to differences in signal strength by adding noise to the cross-item decoder to match it with the regular decoder in terms of mean and standard deviation (Materials and Methods). Importantly, the interference effects with RT remained significant after noise matching (experiment 1: t_19_ = −2.818, p = 0.011, BF_H1_ = 4.741; experiment 2: t_29_ = −1.889, p = 0.034, BF_H1_ = 1.786; pooled data: t_42_ = −3.158, p = 0.001, BF_H1_ = 22.922), emphasising that differences in signal strength do not explain away differences between functional states (Fig. 7A, B).

**Figure 7:**
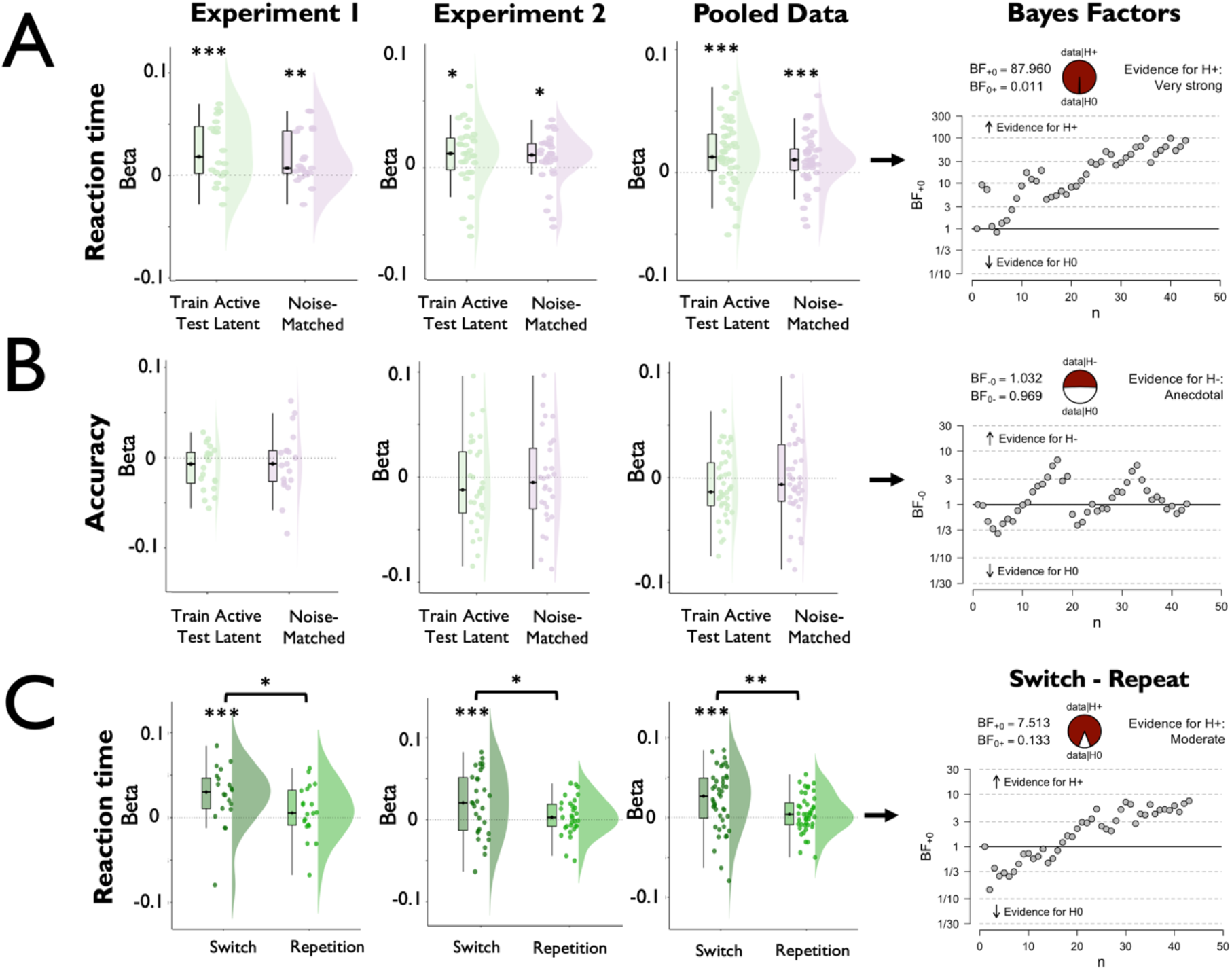
Regression of task performance based on trial-wise variance in the decoding strength of a cross-item decoder that was trained on the data sorted by the cued WM item and tested on the data sorted by the uncued WM item. Results are shown separately for experiment 1, experiment 2, and the pooled data across experiments. The cross-item decoder reliably predicted slower reaction time (**A**, green plots), but was unrelated to accuracy (**B**, green plots)). Notably, the RT effect remained significant after adding noise to cross-item decoder output to match it in signal strength with the regular decoder of the uncued item by adding noise to the decoder output (purple plots; see Materials and Methods). **C** Reaction time effects of the cross-item decoder were only observed on trials that required a priority shift between WM items relative to the previous trial (switch trials), but not on trials in which the item priority was repeated (repetition trials). Plots display the distribution of regression weights across participants. Boxplots: centre lines indicate the median, the box outlines indicate the 25^th^ and 75^th^ percentiles, and the whiskers indicate 1.5 times the interquartile range. Plots on the right display sequential Bayes Factors for the respective regression weights (based on the pooled data set of the cross-item decoder).

We reasoned that instances of cross-item interference might be expressed most strongly on trials demanding priority shifts between items, and therefore repeated the foregoing analysis separately for trials incurring priority shifts, relative to the previous trial (switch trials), and trials that did not (repetition trials). Indeed, the negative prediction of log-RT was significant only on switch trials (experiment 1: t_19_ = 3.259, p = 0.002, BF_H1_ = 21.499; experiment 2: t_29_ = 2.774, p = 0.005, BF = 9.270; pooled data: t_42_ = 3.790, p < 0.001, BF_H1_ = 116.313), but not on repetition trials (experiment 1: t_19_ = 1.127, p = 0.137, BF_H0_ = 1.450; experiment 2: t_29_ = 0.961, p = 0.172, BF_H0_ = 2.060; pooled data: t_42_ = 1.347, p = 0.093, BF_H0_ = 1.452), and the difference between trial types was significant (experiment 1: t_19_ = 2.034, p = 0.028, BF = 2.450; experiment 2: t_29_ = 1.966, p = 0.029, BF = 2.028; pooled data: t_42_ = 2.675, p = 0.005, BF_H1_ = 7.513). Collectively, these results show that interference between WM items arises due to lingering neural patterns coding for previously acted upon, but no longer immediately task-relevant, items.

### Drift Diffusion Modelling

We conducted another set of analyses to obtain more detailed insights into the mechanisms by which functionally active WM states influenced decisions in our task and compare two different explanations for the link between decoding strength and performance (see Figure 8 for illustration): firstly, the strength of the functionally active state may determine the ease with which sensory input is interpreted.^31,32^ This would be the case if the functionally active WM item acts as a matched filter that directly feeds in the accumulation of decision-related evidence (matched filter hypothesis). Secondly, the strength of the functionally active state may determine the ease with which the item can be retrieved for decision making,^33^ thus providing a head-start for decisions on trials with a strong representation of the cued item (retrieval head-start hypothesis).

**Figure 8:**
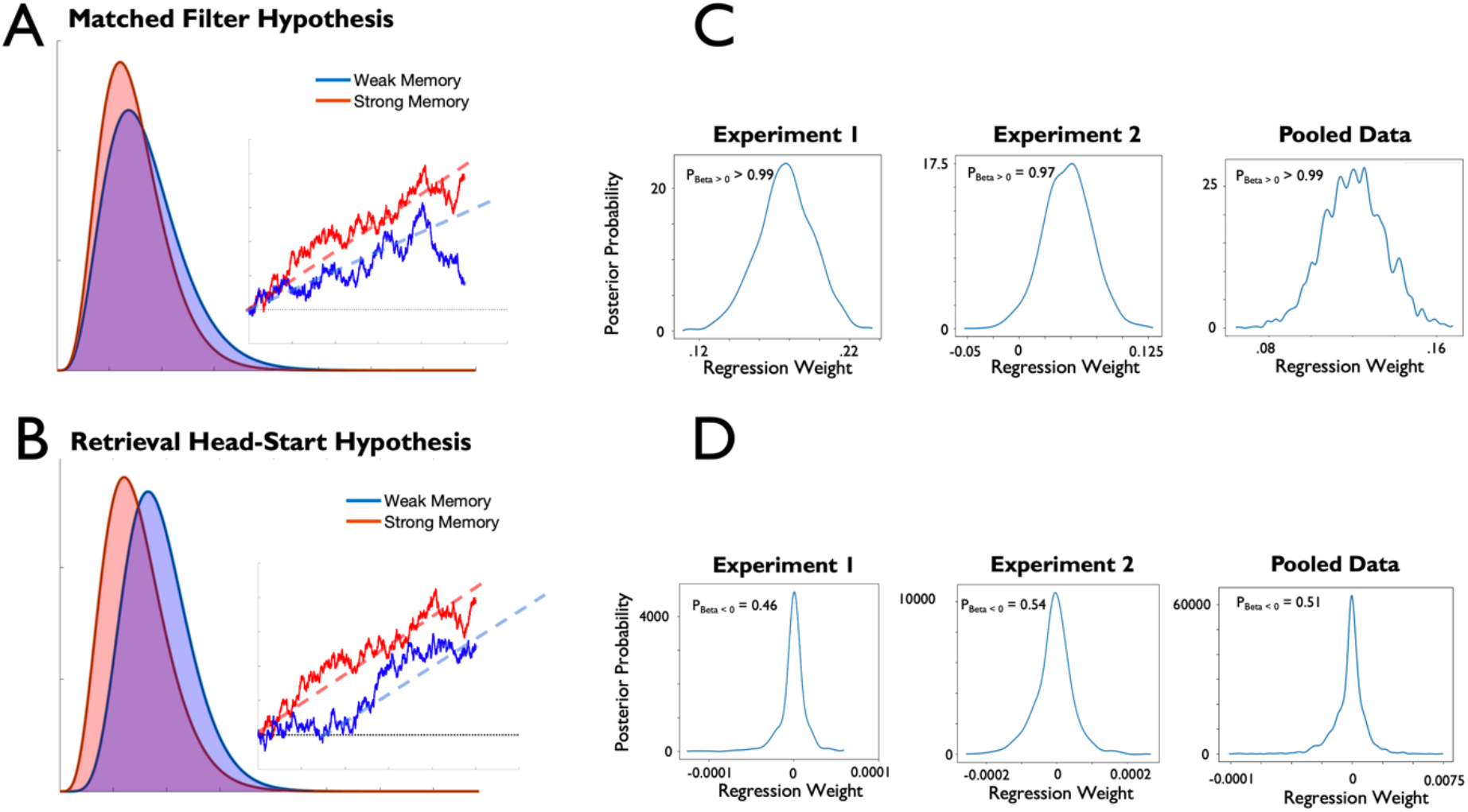
Drift diffusion modelling of task performance as a function of decoding strength of the active WM item. **A**, **B**, Illustration of the two hypotheses under investigation that predict that trial-wise variance in decoding of the cued item tracks either changes in drift rate (matched filter hypothesis) or non-decision time (retrieval head-start hypothesis). The plots display simulated RT distributions and exemplary single-trial diffusion patterns under the two accounts. **C**, **D** Results of the DDM regression analysis that predicted trial-wise changes in drift rate (**C**) and non-decision time (**D**) based on trial-wise changes in decoding strength of the cued item. Plots display the posterior distribution of estimated regression weights, resulting from 5,000 iterations from which the initial 1,00 iterations were discarded as burn in. The significance of effects was evaluated by quantifying the amount of the posterior probability mass that was in the predicted direction (positive for drift rate and negative for non-decision time; see Materials and Methods and Results for details). As indicated on the plots, decoding of the cued item reliably predicted changes in drift rate, but it was not associated with changes in non-decision time.

To evaluate these possibilities, we fit a set of drift diffusion models (DDMs) to our behavioural data and related variance in model parameters to variance in decoding. We focused on two parameters of the DDM: drift rate and non-decision time. The *drift rate* reflects the quality with which decision-relevant information is integrated, and scales negatively with categorisation difficulty. The *non-decision time* reflects the time needed for processes that are not directly related to evidence accumulation such as the encoding of a stimulus or the execution of a response. To test the accounts outlined above, we fit a set of linear regression models predicting trial-wise changes in the two decision parameters based on trial-wise changes in decoding strength of the cued memory item (see Materials and Methods). The matched filter hypothesis predicts that stronger decoding is associated with higher drift rates, whereas the retrieval head-start hypothesis predicts that stronger decoding is associated with reduced non-decision time (see Figure 8). In support of the matched filter hypothesis, we observed reliable positive regression weights for predictions of drift rate in both experiments (experiment 1: P_Beta > 0_ = 0.99; experiment 2: P_Beta > 0_ = 0.97; pooled data: P_Beta > 0_ = 0.99), whereas regression weights for the non-decision time were indistinguishable from zero (experiment 1: P_Beta < 0_ = 0.46; experiment 2: P_Beta < 0_ = 0.54, pooled data: P_Beta < 0_ = 0.51).

### Priority shifts and WM precision

In the final analyses, we aimed to further characterise the representational format of functionally latent WM states by testing if shifting priority away from an item would perturb its precision. As outlined above, several models assume that uncued items are degraded versions of cued items, e.g., because they may receive a smaller part of a limited resource for WM maintenance. In contrast, state-based models assume that cued and uncued items differ only with regard to their functional properties but not with regard to their precision. Importantly, the block structure of our task, with multiple sequences of item priority (i.e., cue sequences; Figure 9A), provided a unique window to adjudicate between these models by examining the precision of WM items before and after priority shifts.

**Figure 9:**
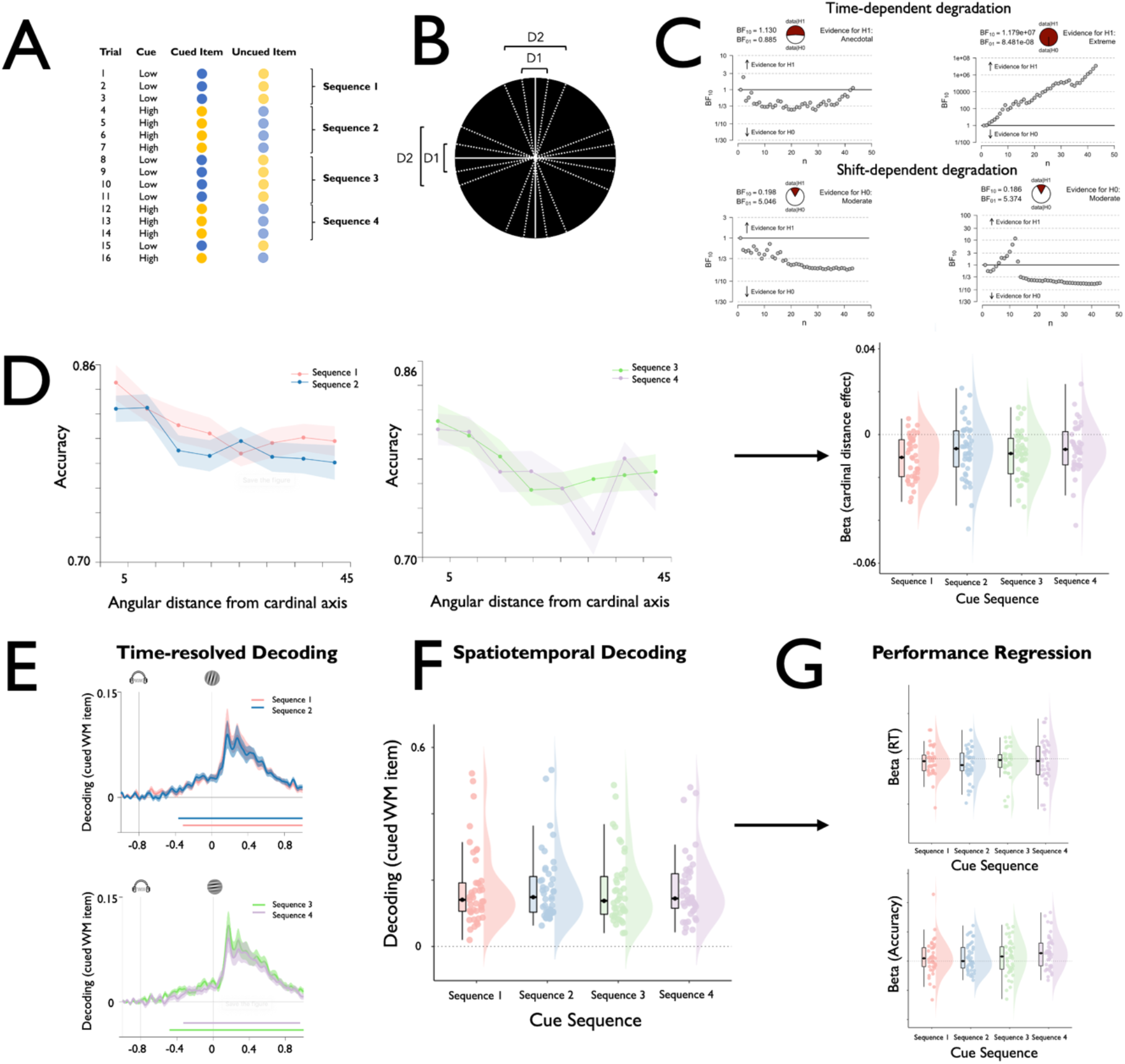
Representational quality of WM items as a function of priority shifts. **A**, Illustration of a hypothetical trial sequence within a task block and the resulting cue sequences that describe periods within the block, wherein an item was used before and after a priority shift. **B,** Illustration of cardinal distances of WM items. For each WM item, we calculated the angular distance to the closest cardinal axis (vertical or horizontal). Note that this visualisation displays a simplified case with only two different distances, whereas our stimulus set contained a total of eight different cardinal distances. **C**, Performance was regressed against the trial number within the current block and the cue sequence within the current block to index time-dependent and shift-dependent degradation of WM items respectively. The plots display sequential Bayes Factors of the respective regression weights for log-RT (left) and accuracy (right). Our results revealed robust effects of time-dependent degradation, but no effects for shift-dependent degradation with Bayes Factors lending support for the null hypotheses. **D**, Performance accuracy was also regressed against the angular distance between the cued WM item and the closest cardinal axis (see B), separately for each cue sequence. The left and the middle plot display mean accuracy for the respective conditions, and the right plot displays the regression weights for each cue sequence. As shown on the plots, we observed robust cardinal distance effects, whereby accuracy decreased with larger distances, but these cardinal bias effects did not differ between cue sequences. We also compared decoding time courses (**E**), spatiotemporal decoding strength (F), and the strength of performance regression (G) of cued WM items between the first four cue sequences. No significant differences were observed. All of these results converge on the notion that priority shifts did not degrade the quality of WM representations Please note that results are shown for the pooled data across both experiments.

**Figure 10:**
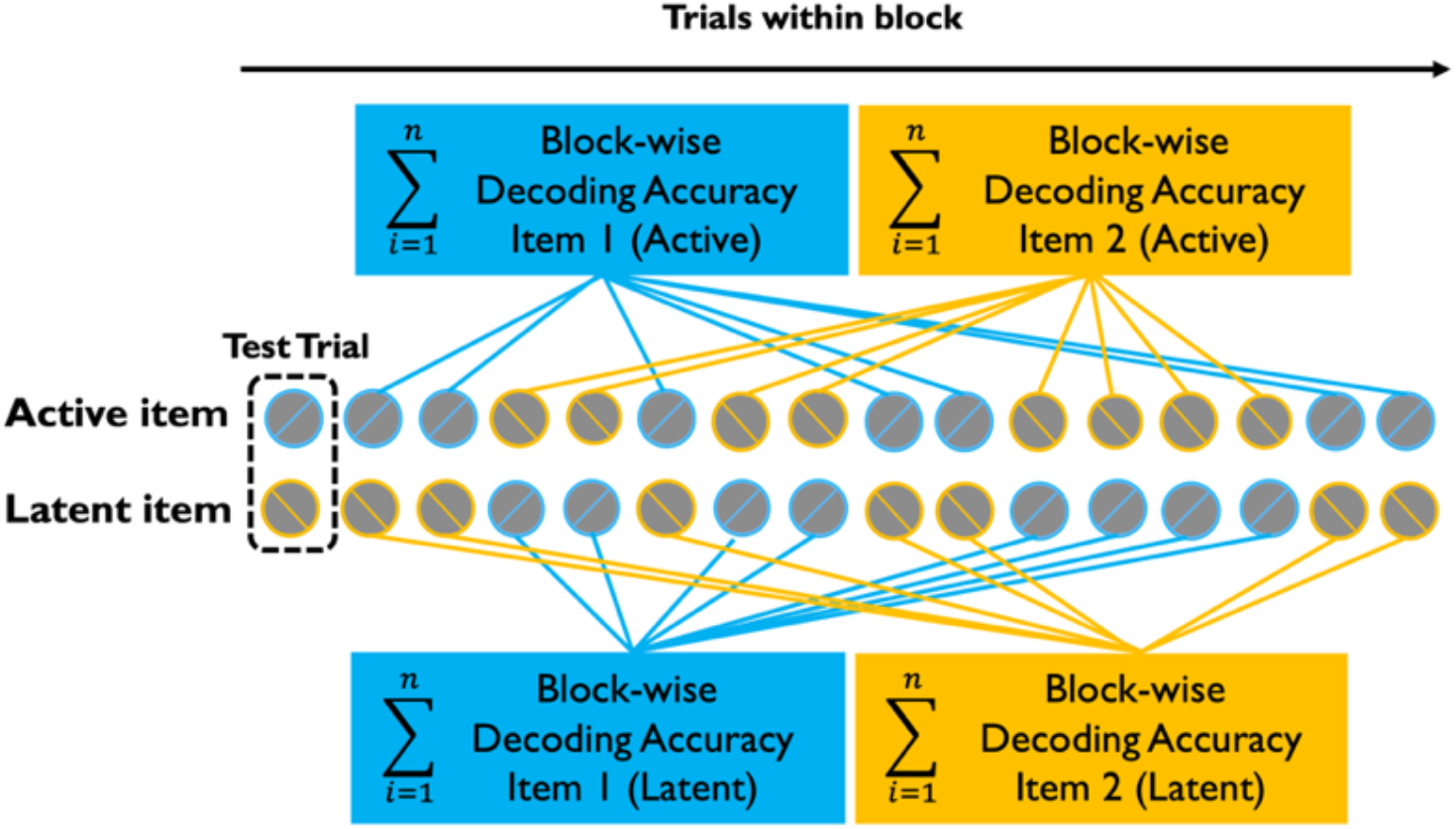
Illustration of the block-wise regression analysis approach. For each of the two WM items within the block, we calculated the average decoding accuracy over all trials on which the respective item task-relevant (top part) and over all the trials in which the respective item was task-irrelevant (bottom part). We then used the block-wise averages to predict trial-wise accuracy and log-RT using the average from all trials except the test trial.

Initially, we used multiple regression to test if the number of priority shifts within a block would predict declines in accuracy beyond the aforementioned time-dependent decline (see Materials and Methods). As shown in Figure 9 C, accuracy declined only as a function of trials within the block (experiment 1: t_19_ = −7.212, p < 0.001, BF_H1_ = 46386.592, experiment 2: t_29_ = 5.382, p < 0.001, BF_H1_ = 4761.628; pooled data: t_42_ = −7.833, p < 0.001, BF_H1_ = 1.179e^7^), but not as a function of priority shifts (experiment 1: t_19_ = −0.799, p = 0.217, BF_H0_ = 2.100; experiment 2: t_29_ = 1.768, p = 0.956, BF_H0_ = 12.961; pooled data: t_42_ = 0.505, p = 0.616, BF_H0_ = 5.374). We next examined if priority shifts would enhance categorical biases, whereby WM items are encoded with respect to the closest cardinal axis^46^ (see Figure 9A and Materials and Methods). Accuracy indeed decreased with larger cardinal distances (experiment 1: t_19_ = −5.801, p < 0.001, BF_H1_ = 3307.866; experiment 2: t_29_ = 6.762, p < 0.001, BF_H1_ = 156142.464, pooled data: t_42_ = 8.475, p < 0.001, BF_H1_ = 8.336e*7), consistent with the presence of categorical biases. Importantly, however, the magnitude of cardinal distance effects did not differ between the first four cue sequences (Figure 9D; all p > 0.441, all BF_H0_ > 3.263), corroborating that priority shifts did not degrade WM precision. Lastly, we compared the EEG-based decoding strength of WM items for the first four cue sequences of each block (see Materials and Methods). Consistent with the foregoing sections, we observed no significant differences between cue sequences in terms of decoding time courses (Figure 9E; all cluster-corrected p > 0.353), spatiotemporal decoding strength (Figure 9F; all p > 0.144, all BF_H0_ > 1.929) or in the link between decoding strength and performance (Figure 9G; all p > .508, all BF_H0_ > 2.354). Altogether, these results show that priority shifts did not degrade memory precision, providing convergent evidence for state-based theories of WM.

## Discussion

In the present study, we compared different forms of WM storage using a task that required maintenance of two items for an extended period of time as well as flexible priority shifts between them. Participants saw the items only during an initial encoding phase and did not receive response feedback throughout the block. Accordingly, while only a single item was task-relevant on each trial, participants had to maintain precise representations of both items to solve the task as a whole. Our results show that cued and uncued items were encoded in qualitatively different functional states that supported performance in different ways: uncued items were encoded in a functionally latent state that was unrelated to performance on the current trial, but tracked memory precision across blocks, suggesting a role in maintaining prospectively relevant memories over intermediate time scales. In contrast, cued items were encoded in a functionally active state that closely tracked dynamic changes in the quality of evidence integration, suggesting a critical role in guiding WM-based decisions over a sub-second time scale. Importantly, functional states were highly flexible and could be dynamically shifted among WM items without degrading their precision.

Our results suggest that the use of WM for guiding behaviour relies on an active reconfiguration of purely mnemonic states into an action-oriented format.^28^ This view aligns with neuropsychological studies documenting dissociations between WM maintenance and use^16,17^, and with studies in healthy humans documenting that cued, but not uncued, WM items can automatically guide attention toward memory-matching stimuli.^18^ Yet previous attempts to characterise WM representations functionally through the use of neuroimaging were hindered by the fact that neural signals coding for uncued items are typically very weak and difficult to measure.^19–22^ In the current study, we were able to recover neural traces of both cued and uncued items from stimulus-evoked EEG signals, permitting us to characterise their functional contributions to WM-based behaviour. Uncued items were encoded in a functionally latent state that did not engage with ongoing processing but was predictive of performance accuracy at a block-wise time scale. This highlights a functional contribution to decision making by providing a storage format that protects memories from decay, while minimising interference with currently prioritised cognition. Such a coding scheme is critical for many complex and time-extended behaviours that incur multiple nested components and priority shifts among them.^35,36^ Understanding how the brain structures cognition over such intermediate time scales will be key for developing more integrative theories of memory-guided behaviour,^37,38^ and we anticipate that such research may reveal contributions to WM maintenance by structures in the medial temporal lobe that have classically been associated with long-term memory.^39^

Cued items were encoded in a functionally active state that predicted dynamic trial-wise changes in performance, especially reaction time. Interestingly, these effects were specifically tied to changes in drift rate, while leaving other component processes reflected in the non-decision time unaffected. This suggests that functionally active WM items act as matched filters that directly transform sensory information into task-dependent decision variables, e.g., by weighting the target-evoked patterns by mnemonic patterns coding for the cued WM item.^31,32,40–42^ At the same time, this result is at odds with proposals that attribute benefits of prioritisation within WM to the facilitation of retrieval processes, which may give decisions a head start, compared to decisions without prioritisation.^33,45^ Notably, the similarity of uncued items to their functionally active state also tracked the amount of interference from those items after priority shifts. This interference was transient and did not cause degradation or forgetting of WM items on subsequent trials, consistent with proposals that the removal and updating of WM contents is an active and time-consuming process^43^, and that failure to complete this process ahead of time leads to interference during target processing.^44^

Importantly, conceiving prioritisation within WM as a transition from functionally latent mnemonic states toward functionally active task-oriented states provides a powerful perspective on other established experimental phenomena such as the effects of retrospective cuing.^47^ These effects have most commonly been explained by assuming a finite resource for WM maintenance that can be continuously distributed among different items.^48^ From this perspective, a more selective focus improves memory for cued items, but at the cost of degraded memory for uncued items. The account we propose offers an alternative explanation, whereby benefits of retrospective cues reflect, at least in part, the reformatting of purely mnemonic states into a prospective action rule that enforces the task-relevant stimulus-response mappings. From this perspective, capacity limitations also concern behavioural readout, in addition to basic maintenance, when assuming that only a single item can be encoded as an action rule at any given moment.^18^ It also implies that selecting an item for guiding behaviour should not necessarily impede the maintenance of other concurrently held items, because once the selected item is encoded in a rule-like state, it is no longer necessary to sustain selective attention to the corresponding mnemonic representation. This could explain previous findings that beneficial effects of retrospective cues can sometimes prevail even when attention is diverted from the cued item,^49,50^ or that they can occur in the absence of reliable performance costs for uncued items.^51,52^ It also aligns with our finding that shifting priority repeatedly between items did not degrade memory precision. The distinction between functionally active and latent WM states also accords well with cases of goal neglect, where WM contents remain accessible, even though they fail to control behaviour.^16^

A central question for future research will be to delineate the precise neurophysiological mechanisms that underpin functional WM states. One possibility is that functionally active and latent WM states directly map onto corresponding activity states: neurally active and ‘activity-silent.’ ^6,53,54^ Our results are consistent with this view, given that spontaneous delay period activity encoded only the cued item, but we could identify traces of both items in the target-evoked EEG signal, consistent with an impulse response uncovering hidden neural states.^12–14,29,30^ However, we note that our current study was not designed to test this specific question but instead focused on dissociating WM states functionally. It is theoretically possible for activity states, or activity-silent states, to support functionally active or latent cognitive states.^55^ The only formal requirement is that their representational format differs qualitatively. This could be achieved by a division of labour between activity-based and activity-silent mechanisms,^53,54^ but also through qualitative differences in the exact patterns,^23,56^ or brain areas,^68^ for respective functional states. In any case, we assert that future research should shift from focusing merely on the presence or absence of decodable memory signals toward characterising their functional properties and the mechanisms that permit their flexible transformation.

One such topic for future inquiry will be to compare the functional states we identified with different mnemonic coding schemes, especially the possibility that cued and uncued items could be represented in inverted activation patterns in the same cortical regions. Two recent fMRI studies that manipulated the priority status of WM items within individual trials observed negative cross-decoding between states,^23,56^ suggestive of opponent coding. At first blush, our findings are inconsistent with this proposal, as we observed positive rather than negative cross-item decoding in our task. There are numerous differences between the two studies and ours, so we can only speculate about the source of this discrepancy. For example, different brain imaging modalities may have captured different aspects of the underlying neural signals with EEG being more sensitive to rapid stimulus-evoked signals and fMRI being more sensitive to tonic and regionally specific signals.^57^ We assume that independent coding schemes for cued and uncued items confer general advantages over inverted schemes, as they permit robust maintenance of both items while minimising interference among them.^55^ However, task demands may alter the optimality of different formats. For instance, inverted representations might become advantageous when simultaneously maintained items are encoded in non-overlapping neural populations, when response contingencies are not explicit before probe onset, or when readout demands require maximal disambiguation between WM items.^58,59^ Therefore, more research will be necessary to establish the boundary conditions of different mnemonic coding schemes.

In conclusion, our results delineate a hierarchical model of WM in which a single item is stored in a qualitatively different format to other concurrently held items. The prioritised format is functionally active and implements a task-relevant transformation of sensory input into decision-evidence, whereas other items are stored in a functionally latent format that does not interact with ongoing processing. Importantly, prioritisation is highly flexible and dynamic, whereby latent states form the basic neural substrate for maintenance and can be used to implement the active representation when needed.

## Materials and Methods

### Participants

20 healthy adults participated in experiment 1 (mean age = 28.1, age range = 18-37, 10 female, 1 left-handed) and 30 healthy adults participated in experiment 2 (mean age = 26.8, age range = 19-41, 14 female, 2 left-handed). Seven participants took part in both experiments. All subjects reported normal or corrected-to-normal vision and received a monetary compensation for participation. The study was approved by, and conducted in accordance with, the guidelines of the Central University Research Ethics Committee of the University of Oxford. For several analyses, we decided to pool the data across both experiments in order to maximise statistical power. In those analyses, we averaged the data from subjects that participated in both experiments across the two sessions, resulting in 43 unique participants for pooled data analyses.

### Apparatus

Stimulus presentation was controlled in Matlab using Psychtoolbox on a 22” monitor with a refresh rate of 100Hz. Unless reported otherwise, stimuli were shown in white font on a grey background (50% contrast, RGB = [127 127 127]) at a distance of approximately 60cm. Responses were given with the left and right index fingers on the “B” and “Y” buttons of a QWERTY keyboard. EEG data were collected using 61 channels that were distributed across the scalp according to the extended 10-20 positioning system. Data were collected at 1000 Hz using a NeuroScan SynAmps RT Amplifier and Curry 7 software. Impedances of all channels were kept below 5 kOhm. In both studies, eye-movements were recorded via electrooculography (EOG) using electrodes places above and below the left eye and to the left of the left eye and to the right of the right eye. In study 2, we additionally recorded eye-movements using a remote infrared eye-tracker (SR Research, Eyelink 1000) sampling both eyes at 1000Hz. We also recorded activity in the first dorsal interosseus muscle of the left and right hand via electromyography (EMG) at 1000Hz.

### Task design

The experimental task required the prolonged maintenance of two items in working memory and flexible priority shifts between those items to guide task-dependent decision making (see Figure 1 for an illustration). Overall, the task was broken into blocks each consisting of 16 trials. At the beginning of each block, participants were shown two orientated bars (6° visual angle in length and 0.25° in width, presented at the same location as subsequent stimuli) in blue (RGB = [25.5 25.5 204]) and yellow (RGB = [204 204 25.5]) colour. These two bars served as memory items for the remainder of the block. For each participant, one colour was associated with a high-pitch tone and the other colour was associated with a low-pitch tone (mapping counterbalanced across participants). The two orientations were drawn randomly from a set of 16 possible orientations (spaced evenly at 11.25° intervals from 2.8125° to 171.5625°) with the sole constraints that the angular distance between the two items could not be identical or exactly orthogonal. Participants could encode the two items for a duration of their choosing and initiated the block via button press. Within the block, each trial started with a presentation of an auditory cue (pure sinusoidal tones, low tone: 440Hz, high tone: 880Hz, duration: 100ms including a 10ms ramp-up and 10ms ramp-down). The cue identity signalled which memory items should be used as a boundary for a forthcoming perceptual decision (cued item), while the other item was maintained merely for later use in the block (uncued item). The cue was followed by a 700ms delay period within which a black fixation dot was presented centrally on the screen (0.15° diameter). Thereafter, a randomly oriented Gabor patch was presented (orientation drawn a set of 16 possible orientations, spaced evenly at 11.25° intervals from 8.4375° to 177.1875°; patches had 6° diameter, 50% contrast, 1.75 cycles/°, random phase, Gaussian envelope with 1.2° SD). Participants were given a maximum time window of 4,000ms to classify the target via button press as being tilted clockwise or counter-clockwise relative to the active memory item. Targets were presented for 100ms and replaced by a fixation dot for the remainder of the response period. In experiment 1, the target was presented centrally on the screen on all trials. In experiment 2, the target was presented laterally at a distance of 6° from the screen centre. The side of target presentation (left vs. right) alternated predictably across blocks. A noise patch (Gaussian smoothed random white noise using a kernel with 0.13° SD, convolved with a Gaussian envelope with 1.2° SD) that matched the target stimulus in luminance, size, contrast, and eccentricity, was presented on the side of the screen at which no target appeared (see Figure 1 for illustration). Clockwise and counter-clockwise decisions were indicated with the right and left index fingers respectively. The response period was followed by a variable intertrial-interval (400-900ms, drawn from a truncated exponential distribution with mean 550ms) until the next trial started with the presentation of a cue. Importantly, participants received feedback about their performance only at the end of each block when their mean accuracy and response time from the preceding block was presented. Accordingly, they had to maintain precise representations of both memory items for the whole duration of the block, as they could not rely on trial-wise feedback to infer the orientation of an item if it was forgotten. Overall, participants completed 128 blocks, resulting in a total of 2048 trials and lasting approximately 2h.

### Behavioural data analyses

Performance accuracy and log-transformed reaction time (log-RT) was analysed using general linear models (GLMs). Initially, we tested to what extent performance was affected by the angular distance between the orientation of the target and the orientation of the cued WM item. We next repeated the analysis but this time testing for an effect of the distance between target orientation and the orientation of the uncued WM item. To visualise these results, we fit the proportion of clockwise choices with a binomial cumulative distribution function using a GLM implemented in R with the ggplot2 and the psyphy packages. We fit another GLM to test for the presence of performance costs as a result of shifting priority within WM by comparing accuracy and RT between trials that incurred a shift in priority (switch trials) and trials that did not (repetition trials). Finally, we tested the stability of performance within and across experimental blocks by predicting accuracy and log-RT based on a variable denoting the trial number within a block (1-16) and another variable denoting the block number within the experiment (1-128).

### Preprocessing

EEG data were initially re-referenced to the average of both mastoids. EEG, EOG, and EMG data were then down-sampled to 250Hz and bandpass filtered, using a high-pass filter of 0.1Hz and a low-pass filter of 45Hz. EEG channels with excessive noise were identified through visual inspection and replaced via interpolation using a weighted average of the surrounding electrodes. The continuous time series data was then divided into epochs, corresponding to the experimental trials starting 200ms before to the onset of the cue and terminating 1800ms after the onset of the target. Each trial was inspected visually for blinks, eye-movements, and non-stereotyped artefacts. Trials were rejected if they contained any of those artefacts during the delay and/or target period. Stereotyped artefacts outside those periods were subsequently removed from the data via independent component analysis. Unless stated otherwise, the data were baseline-corrected for the decoding analyses using the average signal from the time window of 200 to 50 ms before cue onset. Eye-tracking data were down-sampled to 250Hz and we identified and interpolated blinks using spline interpolation and a time window of ±100ms around the event.^61^

### EEG decoding analyses

We conducted a series of multivariate pattern analyses to characterise the neural representations that underpin performance in our task.

### Time-resolved decoding

In a first step, we conducted a *time-resolved decoding analysis* to reveal the time courses at which three variables of our task were explicitly encoded in EEG sensor activity: 1) the orientation of the target stimulus, 2) the orientation of the active and latent memory item, and 3) a decision variable that was calculated as the absolute distance between the orientation of the target and the orientation of the two memory items. To recover parametric information about these variables from EEG sensor activity with high temporal resolution, we computed Mahalanobis distances between the patterns of sensor activity that were evoked by different stimulus orientations and measured the extent to which these distances reflected the underlying circular orientation space (or linear decision variable space). We used a leave-one-block-out cross validation procedure with the data from each block serving as test data once and all the remaining data serving as training data. Training data were subdivided into 16 bins, according to their orientation relative to the test data, and then averaged. Mahalanobis distances between the 16 average patterns in the training data and each test trial data were then computed using the noise covariance from training data. Noise covariance was calculated on the average residual data after subtracting orientation-specific mean activity from each trial with the corresponding orientation. Residuals were averaged over a time window from 0 to 1.8s relative to the onset of the auditory cue (and therefore encompassed the cue, delay, target, and decision phases). The covariance was calculated over the trials-by-sensors matrix of average residuals, using a shrinkage estimator.^62^ This procedure was repeated for every block and for every time point within the trial using 4ms time bins. We normalised the resulting pattern distances by subtracting the mean across the 16 distances for each time point from each sensor’s activity. To simplify interpretation, we also reversed their sign so that positive values reflected pattern similarity rather than distance. This procedure enabled us to compute an orientation tuning curve for each time point and trial, which expresses the extent to which pattern similarity decreases as a function of angular distance. To transform the 16-dimensional tuning curve into a one-dimensional index of decoding accuracy, we computed the cosine vector mean of each tuning curve by rescaling the cosine of the centre of each orientation bin to the range from −180 to 180 and multiplying it with the corresponding pattern similarities. The mean of the resulting 16 values served as index of decoding accuracy with positive values reflecting tuning of the EEG signal for stimulus orientation.^14,22^ Decoding values were smoothed with a Gaussian kernel (SD = 24ms) for visualisation and significance testing. We tested for significance using one-sample t-tests against 0 at each time point, and corrected for multiple comparisons in time via cluster-based permutation testing using 10,000 permutations.^63^ For the decoding of the memory items, cluster correction was applied for the whole trial. In contrast, for the decoding of the target and the decision variable, cluster correction was only applied for the target period (i.e., the 1800ms after target onset), because these variables could by definition only be encoded after the onset of the target stimulus. In keeping with previous work from our group,^14,29,30^ decoding analyses were conducted only within posterior EEG sensors (P7, P5, P3, P1, P2, P4, P6, P8, PO7, PO5, PO3, POz, PO4, PO8, O1, Oz, O2). We selected these channels to be consistent with previous studies and because orientation signals are typically expressed most strongly in posterior regions.^14,32,64,65^ Please note, however, that none of the reported results depend on this channel sub-selection (i.e., all reported effects remain significant when using all 61 channels). Moreover, please note that in experiment 2 decoding analyses were conducted separately for blocks with target presentation on the left and right side, and decoding accuracies were subsequently averaged across sides. The decision variable (absolute difference between memory item and target) followed a discrete uniform distribution between 5.625° and 84.375°, unlike the other variables (memory items and target stimuli), which had a circular distribution. Our decoding approach therefore needed to be adjusted for decoding of the decision variable. Instead of reducing the dimensionality of the tuning curve using a cosine vector mean, we calculated the linear slope of the tuning curve.

### Spatiotemporal decoding from stimulus evoked EEG signals

In a second step, we employed a complementary analysis approach to decode the same task variables from longer time-windows of stimulus-evoked brain activity. This approach exploits the dynamic temporal structure of event-related potentials by pooling multivariate information in time, thus capturing not only information encoded in spatial activation patterns, but also information that is encoded in the temporal unfolding of these spatial patterns. A previous study from our group has shown that such pooling over adjacent time points within trials can increase decoding sensitivity at the expense of temporal precision.^29^ Consequently, the two decoding approaches capture complementary aspects of neural representations and should be considered mutually informative. In keeping with the previous study, we combined EEG data from the 17 posterior channels within a time window of 100-400ms after target onset, thus treating single trials as discrete events. The mean activity at each sensor and time point was removed to normalise voltage fluctuations and isolate dynamic, stimulus-evoked brain states from brain activity that is stable across the chosen time period. To avoid having more data features (number of sensors X number of time points in the training data) than data samples (number of trials in the training set), we also down-sampled the signal at each sensor by a factor of 10 (i.e., from 250Hz to 25Hz). Following this preparation, we performed decoding analyses via Mahalanobis distances using the same procedure as detailed above. The 4D approach was used to decode the same task variables (target stimulus, memory items, decision variable, decision category), yielding a single vector of decoding accuracies for each subject and task variable. We evaluated the significance of results via one-sample t-test against chance level (one-tailed, testing for above-chance decoding).

### Regression analyses

We conducted a series of regression analyses to establish the behavioural relevance of the decoded memory signals and test our hypothesis that active and latent memory items are encoded in qualitatively different states. In a first step, we regressed log-transformed reaction time (log-RT) and accuracy against the trial-wise decoding strength of cued and uncued WM items. That is, for each trial and item, we calculated the decoding strength using the spatiotemporal approach described above, and normalised these scores by subtracting the block average to isolate across-trial fluctuations from more sustained changes in decoding. We then used these normalised scores to predict trial-wise variance in performance with general linear models serving to predict log-RT and logistic regression models serving to predict accuracy. We also included regressors for the block number and the trial within the block to account for general changes in performance across and within blocks. Given that we were interested in potential null effects (i.e., absence of links between trial-wise variance in performance and trialwise variance in the decoding strength of uncued items), we analysed the resulting regression weights through a combination of frequentist and Bayesian statistics. Initially, we tested if regression weights were significantly different from zero using one-sample t-tests and also calculated Bayes Factors from Bayesian one-sample t-tests to quantify the evidence in favour of the null hypothesis and the alternative hypothesis. Based on our hypotheses, we used one-tailed tests in the direction of facilitation (i.e., positive regression weights for accuracy and negative weights for log-RT) for cued items and in the direction of interference (i.e., negative weights for accuracy and positive weights for log-RT) for the uncued item These analyses were implemented using JASP software using default priors for the Bayesian analyses.^67^ We also compared regression weights between cued and uncued items using paired t-tests (one-tailed, testing for greater facilitation with cued items) and calculated Bayes Factors resulting from a Bayesian paired-samples t-test.

In a second step, we regressed log-RT and accuracy against the average block-wise decoding strength of each memory item. That is, we predicted performance on each trial (test trial) based on the average decoding strength across all remaining trials within the same block on which the respective memory item was cued or based on the average decoding strength across all trials on which the respective memory item was uncued (see Figure 9 for illustration). Analogously to the trial-wise analyses, we calculated frequentist and Bayesian statistics to analyse the resulting regression weights.

Finally, we repeated the trial-wise analysis with the output from cross-item decoding analyses by training the pattern classifier on the data sorted by the cued item and tested on the data sorted by the uncued item. This analysis aimed to test if interference from uncued items scales with the extent to which uncued items are encoded in neural patterns that resemble the items functionally active state. Initially, the cross-item decoders were tested against chance level using one-sample t-tests (one-tailed, testing for above-chance decoding), and compared with the regular decoder of the uncued item using paired samples t-tests and Bayes Factors (two-tailed, based on absence of a priori predictions). Thereafter, the regression weights of the cross-item decoder were tested against zero using one-sample-t-tests and Bayes Factors (one-tailed, testing for interference effects). We repeated this analysis separately for trials that demanded a priority shift between WM items (switch trials), relative to the previous trial, and trials that did not (repetition trials). Regression weights were compared between trial types via paired samples t-tests (one-tailed, testing for greater interference effects on switch trials). Finally, we also conducted a control analysis to match the cross-item decoder and the regular decoder of the uncued item in terms of mean and standard deviation. To this end, we first mean-centred the two vectors that denoted trial-wise scores of each decoder. Mean-centred vectors were then scaled by the standard deviation of the cross-item decoding vector. We then calculated the standard deviation of the noise that was required to match the scaled cross-item decoder with the scaled regular decoder of the uncued item and added random noise with the estimated properties to the output of the cross-item decoder. The noise-matched decoder was then used to predict performance using the method as described above for trial-wise regression analyses. This procedure was repeated with 1,000 noise injections and regression weights were averaged across all iterations.

### Drift Diffusion Modelling

In the next set of analyses, we aimed to obtain more detailed insights into the way memory strength influenced WM-guided decisions in our task. To this end, we fit a set of drift diffusion models (DDMs) to our behavioural data and regressed variance in model parameters to variance in decoding. DDMs characterize decisions as the accumulation of noisy evidence between two competing options, whereby one option is chosen once a sufficient amount of evidence has been accumulated.^34,45^ DDMs decompose performance in two-alternative choice tasks into latent decision parameters based on RT distributions and choice probabilities. The most parsimonious version of the DDM has three parameters: drift rate, non-decision time, and decision threshold. Firstly, the *drift rate* reflects the quality or strength of decision-relevant information and scales negatively with categorisation difficulty (i.e., lower drift rate with more difficult discriminations). Secondly, the *non-decision time* is thought to reflect the time needed for processes that are not directly related to evidence accumulation such as the encoding of a stimulus or the execution of a response. Finally, the *decision threshold* (or boundary separation) reflects the amount of evidence that is needed to commit to a behavioural choice. This parameter is thought to be under the strategic control and to regulate speed-accuracy trade-offs (e.g., higher thresholds will result in slower but more accurate responses).

After observing that trial-wise variance in decoding of the active, but latent, WM item tracked variance in task performance (see Results section), the central aim of this analysis was to identify which decision parameter could account for this benefit. This enabled us to compare two hypotheses: (1) the matched filter hypothesis and (2) the retrieval head-start hypothesis. Under the matched filter hypothesis, variance in decoding strength should track the ease with which sensory input is interpreted and transformed into a decision variable. This view predicts that variance in decoding across trials should be positively associated with variance in drift rate (see Figure 8). By contrast, under the retrieval head-start hypothesis, variance in decoding strength should track the accessibility of the active WM item for decision making. This view predicts that variance in decoding across trials should be negatively associated with variance in non-decision time (see Figure 8).

To test these accounts, we conducted a regression analysis using a hierarchical Bayesian version of the DDM implemented in the HDDM python toolbox under version 0.6.^66^ HDDM applies Markov Chain Monte Carlo sampling methods to estimate posterior distributions over DDM parameters. We fit a set of linear regression models using the patsy library to determine the relation between trial-wise fluctuation in decoding strength for both cued and uncued items and trial-wise fluctuation in drift rate and non-decision time respectively. We used performance accuracy (correct vs. incorrect) as response bounds rather than response identity (clockwise vs. counter-clockwise) because we aimed to establish links between decoding strength and performance, and had no reason to suspect a response bias. Separate analyses were run for drift rate and non-decision time. Each analysis contained 5000 samples from which the first 1000 were discarded as burn-in. We evaluated results statistically via the posterior distribution of the estimated regression weights. Effects were considered significant if at least 95% of the posterior probability mass was above or below zero, and in the predicted direction (positive for drift rate and negative for non-decision time; see explanation above).

### Effects of priority shifts on memory precision

In the final set of analyses, we aimed to characterise the format of functionally latent WM states by testing if shifting priority away from a WM item would degrade its precision. To this end, we conducted a series of behavioural and EEG analyses. Initially, we performed another regression analysis to test if the number of priority shifts that had preceded a given trial within a block would predict a decline in performance above and beyond a purely time-dependent decline (as measured by the trial number within the blocks). We predicted accuracy and log-RT using a GLM that contained predictor variables for the trial number within the current block (block trial) and the cue sequence (i.e., the number of cue switches preceding the current trial within the; see Fig 9A). The block trial variable indexes time-dependent degradation, whereas the cue sequence variable indexes shift-dependent degradation. As for the previous analyses, we evaluated regression weights of each predictor variable using one-sample t-tests and Bayes Factors from Bayesian one-sample-t-tests.

In a second step, we tested whether shifting priority away from an item would increase categorical biases, whereby items are encoded with respect to their closest cardinal boundary. We therefore calculated a variable reflecting the angular distance between the cued WM item on each trial and closest cardinal axis (vertical or horizontal). This variable was used as a predictor for accuracy and log-RT, separately for each cue sequence. Regression weights were again evaluated using one-sample t-test and Bayes Factors; and compared between cue sequences using paired samples t-tests and Bayes Factors. Paired samples t-tests were one-tailed testing for shift-dependent degradation (i.e., larger effects of cardinal distance with later cue sequences).

Finally, we also tested effects of priority shifts on WM representations using EEG-based decoding strength as an index of WM precision. We therefore divided decoder outputs based on the first four cue sequence within each block (see Figure 9A). To test for differences in decoding time course, we compared the time-resolved decoding strength between the four cue sequences using the cluster correction described above. We also compared the spatiotemporal decoding strength (see description above) between the first four cue sequences using paired-samples t-test and Bayes Factors (one-tailed, testing for shift-dependent degradation). Lastly, we also conducted regression analyses, predicting log-RT and accuracy from spatiotemporal decoding strength, separately for each cue sequence. As for the previous analyses, regression weights were tested against chance using one-sample t-tests and Bayes Factors, and compared using paired-sample t-tests and Bayes Factors (one-tailed, testing for decreases in performance prediction with larger cue sequences).

## Acknowledgement

This research was supported by the Wellcome Trust (award 210849/Z/18/Z to PSMK and award 201409/Z/16/Z/ to NEM), Linacre College Oxford (PSMK), University College Oxford (NEM), the Lucy Halsall Foundation (PSMK), the Research Foundation Flanders (award 12R8817N to PSMK), the Biotechnology and Biological Sciences Research Council (award BB/M010732/1 to MGS), the James S McDonnel Foundation (award 220020405 to MGS), and the NIHR Oxford Health Biomedical Research Centre. The Wellcome Centre for Integrative Neuroimaging is supported by core funding from the Wellcome Trust (203139/Z/16/Z). The authors would like to thank Dan Bang and Annika Boldt for advice on the drift diffusion modelling, Michael Wolff for support with Figure 2, and Thomas Christophel, John Duncan, Sanjay Manohar, and Darinka Trübutschek for comments on previous versions of the manuscript.

## Supplementary Material

**Supplementary Figure 1:**
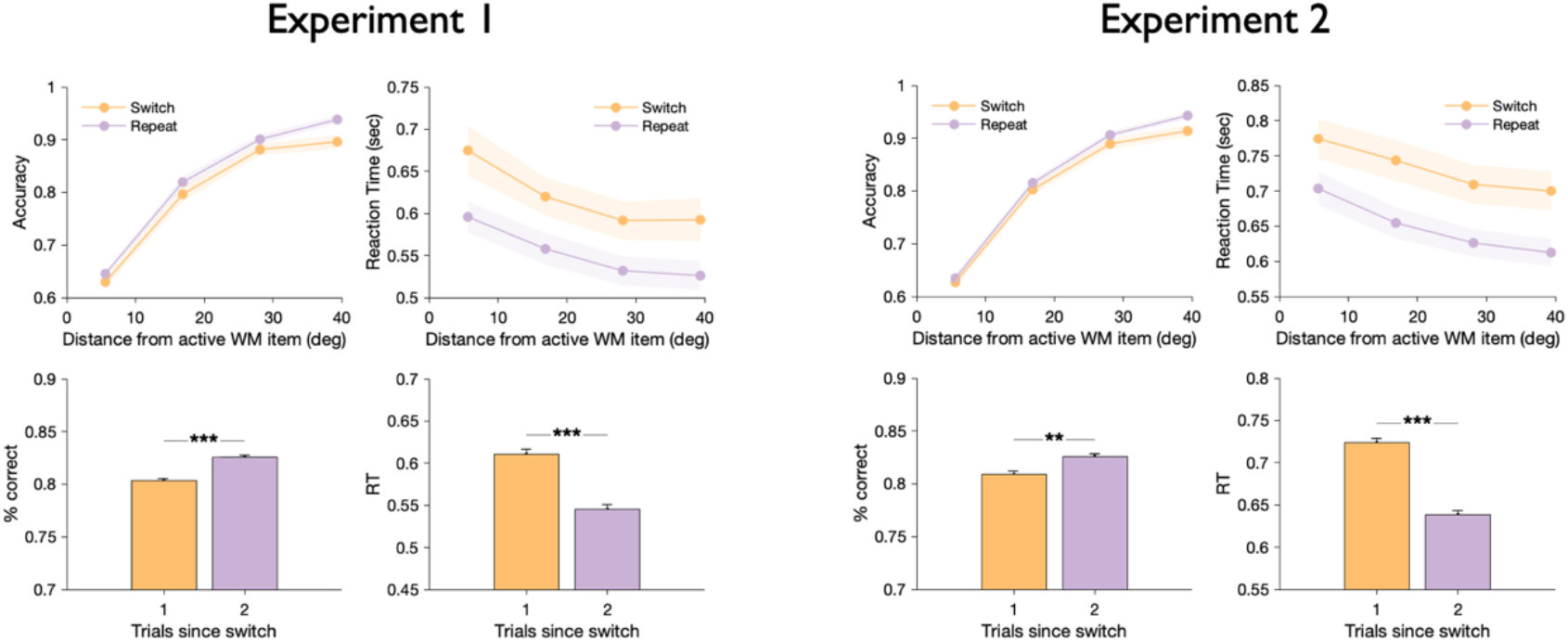
Behavioural effects of priority shifts between WM items were observed in experiment 1 (left plots) and experiment 2 (right plots) whereby performance was less accurate and slower on the first trial after a switch (bottom panels). These effects were observed across all angular distances between the target orientation and the orientation of the active WM item (upper plots). ** p < 0.01, *** p < 0.005. Shadings and error bars indicate standard error of the means.

**Supplementary Figure 2:**
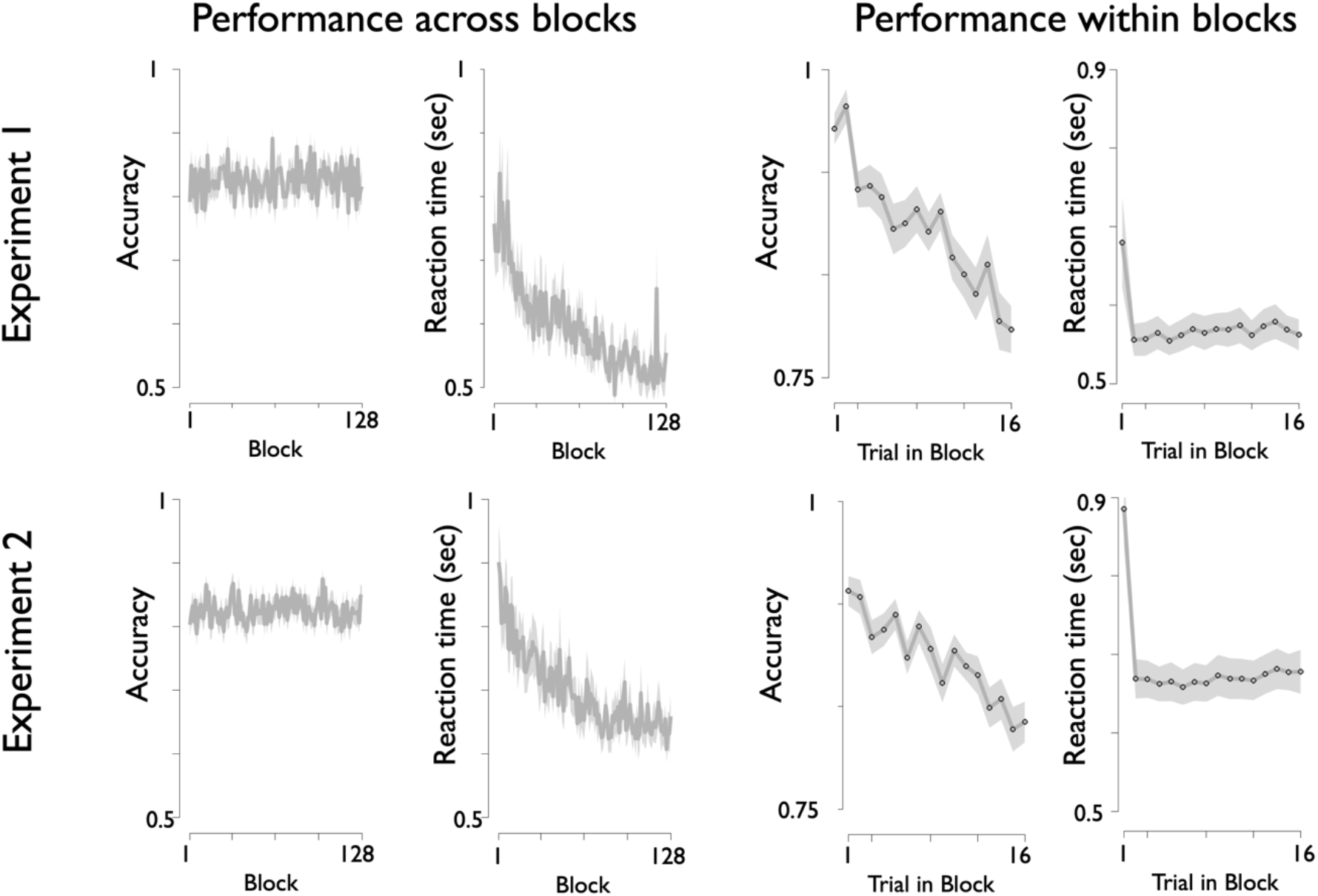
Performance stability across and within experimental blocks for experiment 1 (top row) and experiment 2 (bottom row). Overall, participants completed 128 blocks, each containing 16 trials. Plots on the left display as a function of block number. While accuracy was stable across blocks, RT continuously declined with increasing block numbers. Conversely, within blocks, accuracy continuously declined, whereas RT was stable with the exception of the first trial, which exhibited longer RT than subsequent trials. Shadings indicate standard error of the means.

**Supplementary Figure 3:**
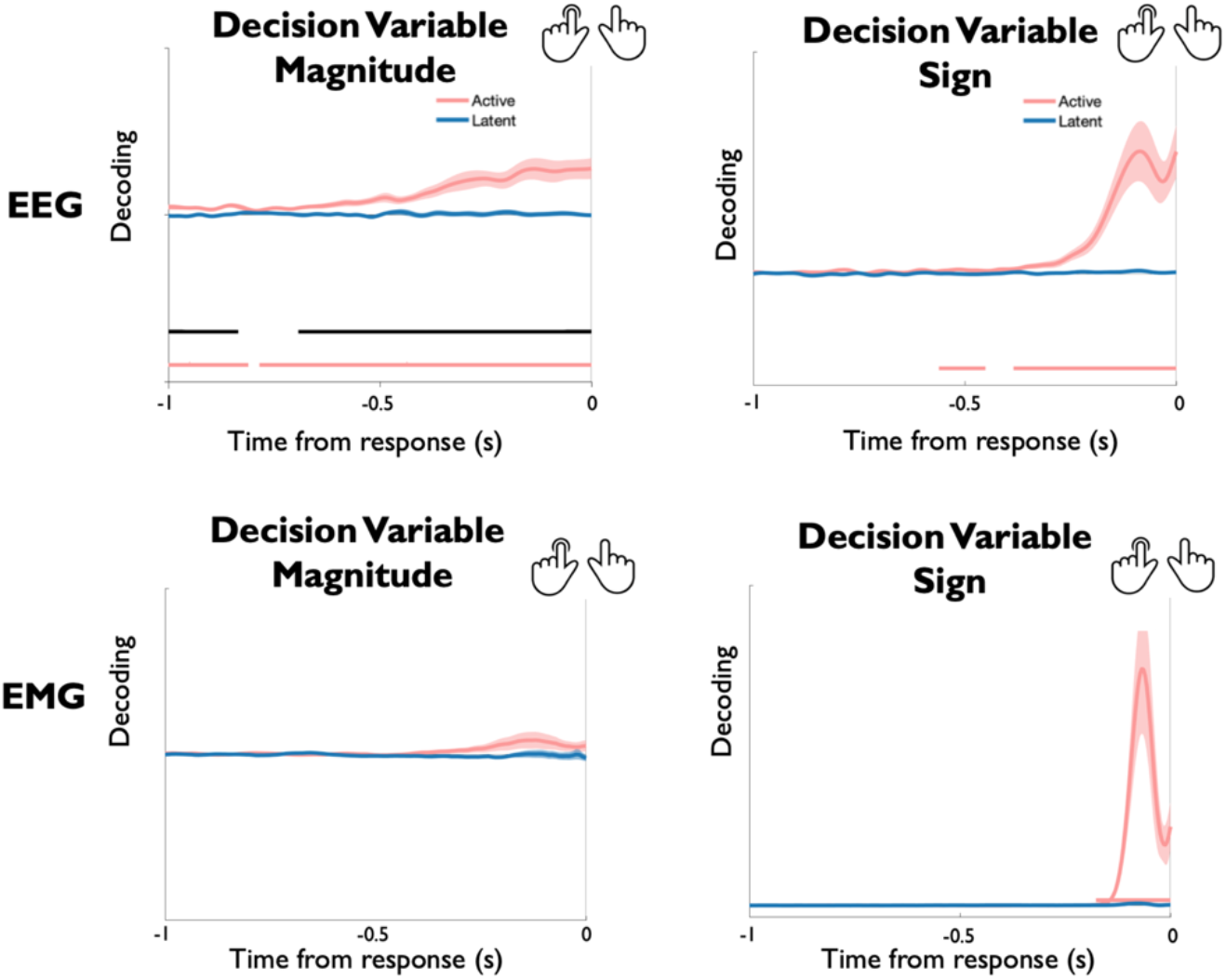
Response-locked decoding of decision variable magnitude (i.e., the absolute angular distance between target orientation and WM item orientation -- equivalent to the decision variable in the main article) and decision variable sign (i.e., a binary variable indicating whether the target was oriented clockwise or counter-clockwise relative to the WM item – equivalent to the behavioural choice). Decoding was conducted separately using the EEG channels (upper plots) and the EMG channels (lower plots) and shown separately for experiment 1 (left plots) and experiment 2 (right plots). The decision variable magnitude signal could only be decoded from EEG channels and ramped up gradually prior to overt responses. In contrast, the decision variable sign could be decoded from EEG and EMG activity and exhibited a sharp signal increase immediately preceding overt responses. These results demonstrate that the decision variable magnitude signal does not index processes of response execution.

**Supplementary Figure 4:**
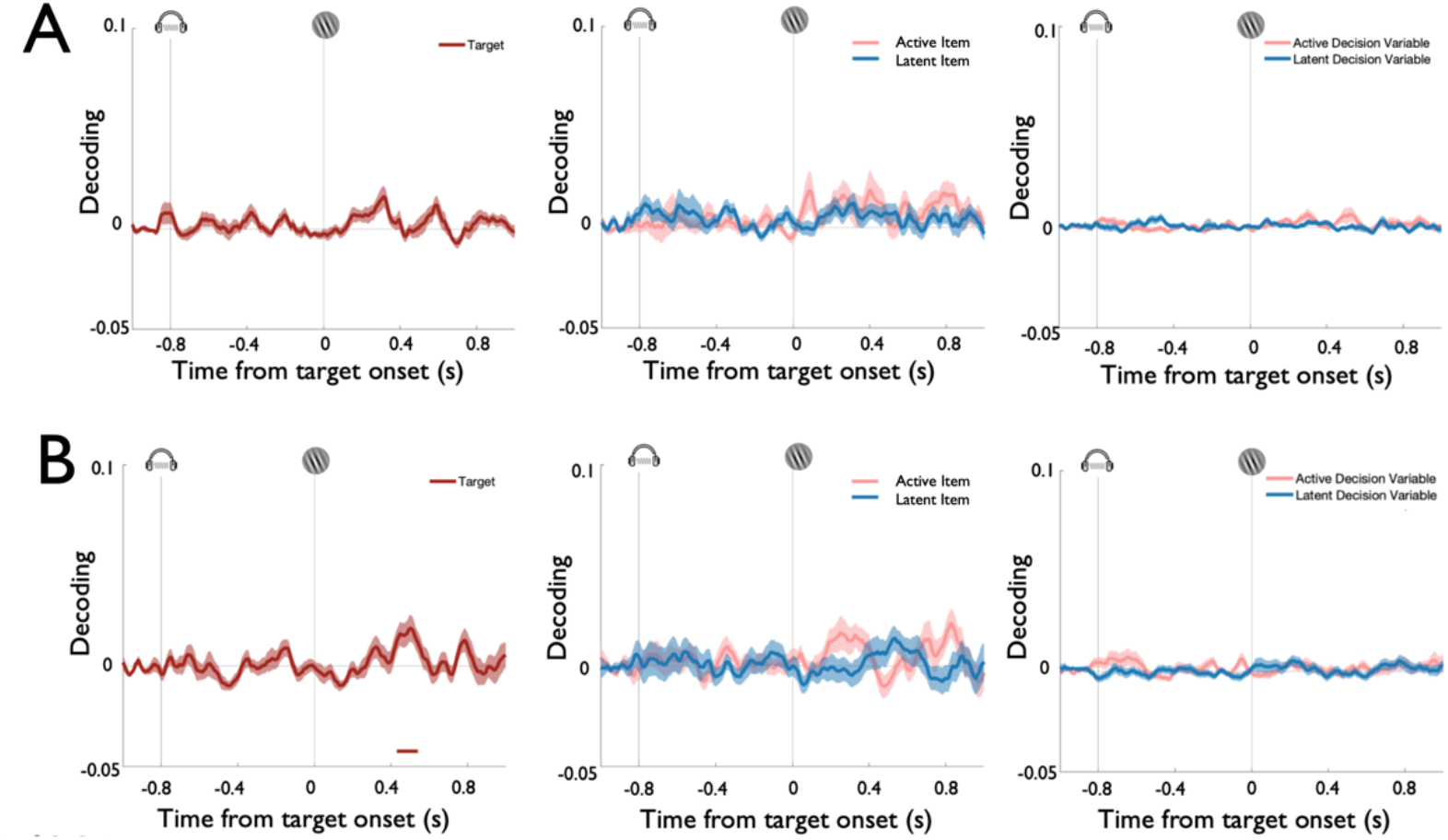
Time-resolved decoding of task variables using EOG channels in experiment 1 (A) and the high-resolution eye-tracker in experiment 2 (B). These data show that eye-channels did not contain reliable information about the task variables that were decoded from EEG sensor patterns. Each plot displays time-resolved decoding results from 200ms before cue onset until 800ms after target onset. Coloured lines represent cluster-corrected time-periods within which decoding was significantly greater than chance. Shading indicates standard error of the mean.

**Supplementary Figure 5:**
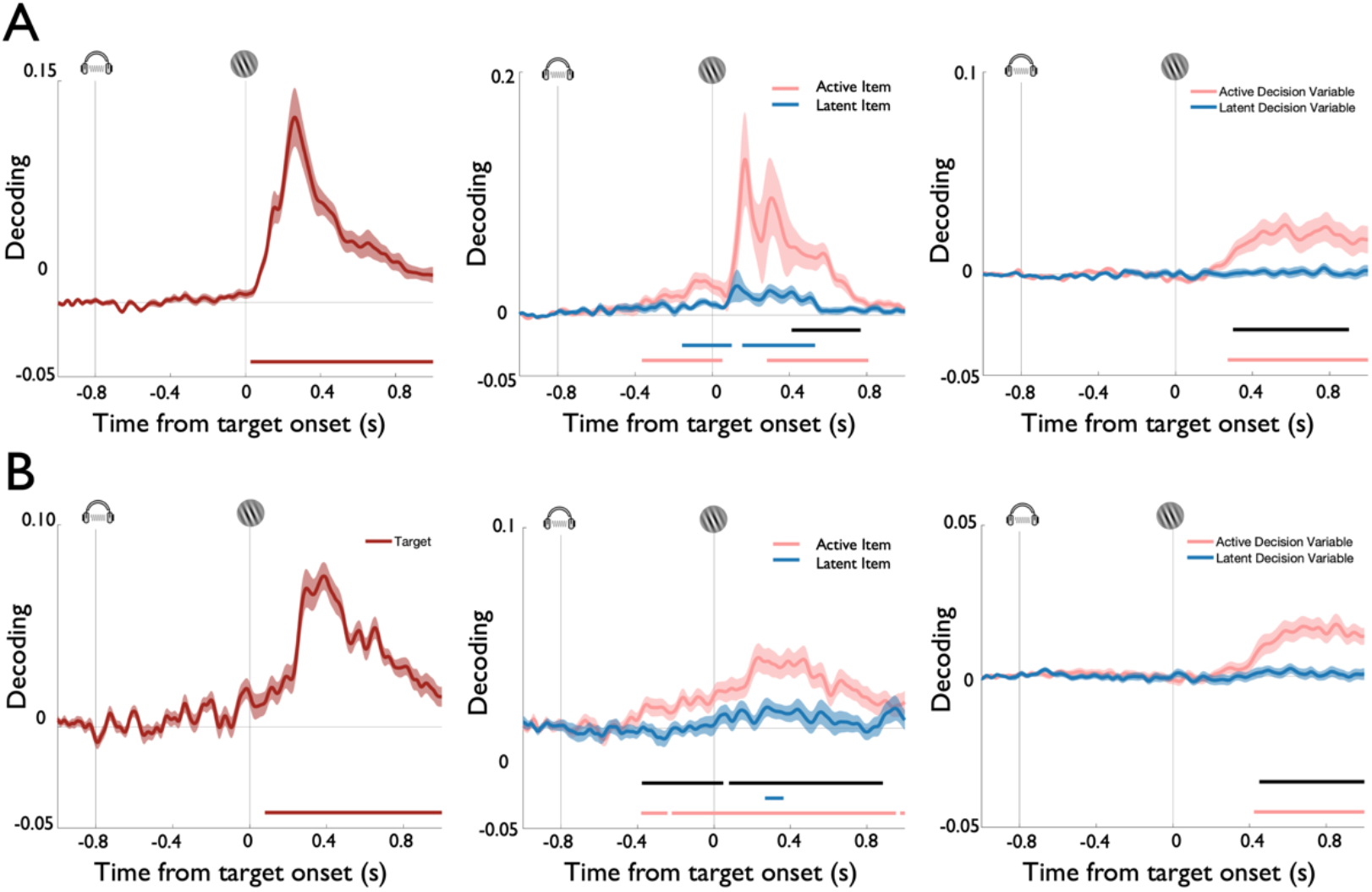
To rule out that the signals decoded from EEG activity were not caused by subtle eye-movements we also regressed decoding time series obtained with the eye-channels against the respective time series obtained with the EEG. Plots indicate the residual EEG signal after regressing out variance that could be explained by eye-channel signals in experiment 1 (A) and experiment 2 (B). No qualitative changes were observed compared to the original EEG decoding analysis (see Figure 3 in the main text), corroborating that eye-movements did not explain the observed decoding results.

**Supplementary Figure 6:**
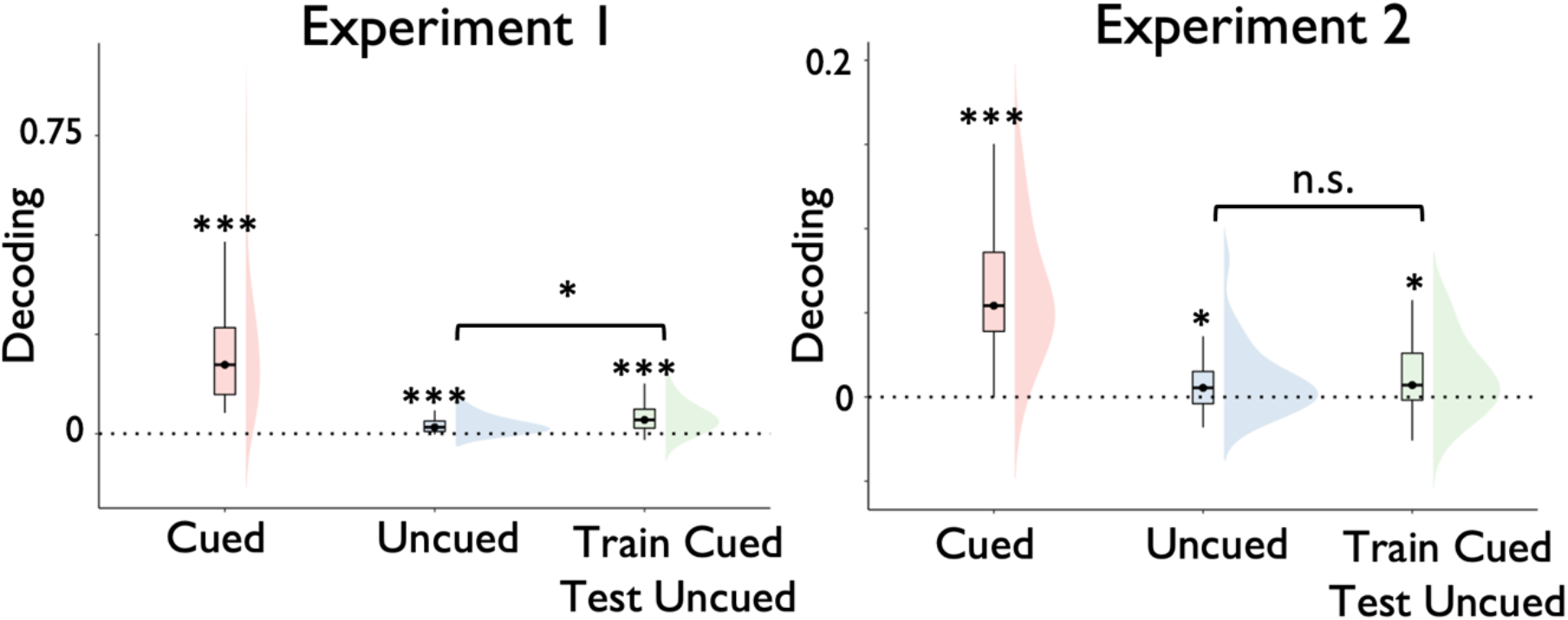
Spatiotemporal decoding results shown for the regular decoders of the cued and uncued WM items and a cross-item decoder that was trained on the data sorted by the cued item and tested on the data sorted by the uncued item. The cross-item decoder classified data with above-chance accuracy in both experiments. The decoding strength was enhanced, relative to the regular decoder of the uncued item, in experiment 1, but not in experiment 2.

